# Trajectories of genetic correlations in populations under selection: from theory to a case-study

**DOI:** 10.1101/2025.03.13.643026

**Authors:** Beatriz C.D. Cuyabano, Mariana R. Motta, Jeremie Vandenplas, Nancy L. Garcia, Fatima Shokor, Pascal Croiseau, Didier Boichard, Sophie Aguerre, Sophie Mattalia

## Abstract

**Background:** Breeding programs select for multiple commercial traits, aiming to achieve genetic progress for all. Often, selection is based on a selection index, *i*.*e*. a linear combination of traits with weights defined by, among other information, the genetic correlation between traits. These correlations are typically estimated as a static parameter, and assumed equal to all individuals and generations. While research on the consequences of selection to genetic variances (Bulmer effect) is widely available, only a few studies focused on the consequences of selection to genetic correlations. Our study extended the already existing inferences about how selection affects genetic variances, to how multi-trait selection affects genetic correlations. In order to further our understanding of genetic correlations, we also proposed an alternative method to calculate genetic correlations between traits at the individual level, called by us as individualized sire genetic correlation (iSGC), obtained through the estimated breeding values (EBV) from evaluated daughters. Lastly, a case-study was performed on thirty years of data from the French Holstein dairy cattle population, for five traits studied pairwise: milk and protein yield, milking speed, somatic cell score, and cow conception rate.

**Results:** Theory revealed that multi-trait selection leads to an attenuation (decrease) of positive genetic correlations, with potential to revert them to negative values, if initially low. Uncorrelated traits will become negatively correlated, and negative genetic correlations will be either intensified or attenuated (decrease or increase, respectively), depending on selection intensity, weights applied to the selection index, and the initial genetic correlation.

**Conclusion:** Both theory and empirical results on real data confirm that selection does change the genetic correlation between traits in a population under selection. Moreover, empirical trajectories of the iSGC were in better agreement with the theory, than trajectories of populational genetic correlations. The iSGC searches for individual-specific patterns of correlations, and since it is measured on sires through the EBV of their daughters, it also considers the recombination of the genetic background. Along with the fact that trajectories of iSGC were in better agreement with theory, we believe it to be a potentially less biased measure of genetic correlations between traits.

## Background

The consequences of selection in genetic variance is a topic that has been widely studied in quantitative genetics, and well described as the Bulmer effect [1, 2], which details the loss of genetic variance driven by selection. [3] further described the changes in genetic variances in populations under selection, as a function of both selection and limited population sizes, since the loss of genetic variance arises due to the build-up of co-ancestry, resulting in inbreeding and random genetic drift [2–4]. Moreover, the current availability of large-scale phenotypic records throughout decades for breeding populations has opened the door to transcend longitudinal studies with respect to variance parameters from the theoretical framework, to studies that aim to verify whether the realized trends in genetic variances are indeed in agreement to the expected, given the goals of a breeding program [5–7].

Most studies about the effects of selection on the genetic parameters have focused in describing and understanding the longitudinal trajectories of genetic variances from a single-trait perspective [1, 3, 6]. It is however, reasonable to expect that the Bulmer effect and random drift have an impact not only in the genetic variances of traits, but also on the genetic correlations between heritable traits involved in a breeding program.

Only a few studies have focused on the consequences of multi-trait selection to both the genetic variances and covariances (thus, correlations) between traits [3, 8–10]. Furthermore, these studies were limited to a few specific scenarios of populations under selection for one or maximum two traits. In plant breeding, studies have also shown an interest in predicting genetic correlations between traits under selection, however specifically for the commercial F1 hybrids of a crossbreeding scheme [11, 12], and by deploying *in silico* simulations assisted by deterministic equations [13].

Evolutionary studies with a genomic approach have more recently turned their attention to understanding the trends of genetic correlations as well [14, 15], however with an evolutionary perspective, seeking to better understand the progress of multiple adaptation and fitness traits. The goals of these more recent evolutionary studies were similar to previous ones, pre-genomics, in this field of research [16, 17], also with a focus in understanding the role played by pleiotropic mutations [18].

In fact, when it comes to the interest of genetic evaluations where the objective lies in predicting breeding values for selection candidates, changes in variance parameters due to selection can be disregarded without compromising the final rank of individual’s genetic merit, a principle known as *ignorable selection* [19–21].

However, when redesigning breeding goals, it is of paramount importance that (co)variance parameters are adequately described and estimated at the current stage of the selected population, so that the expected medium to long-term genetic progress in achieved for all objective traits. When it comes to building an individual’s selection index based on multiple traits, an extra special attention must be drawn to the relevance of correct inputs for the genetic correlations, as incorrect inputs may compromise particularly the expected trade-off between negatively correlated objective traits.

Moreover, the longitudinal study of genetic (co)variances may enable inferences for traits that are not routinely or currently measured, but that may be included in future breeding goals. Such traits, when correlated to those already featuring in a selection index (SI), are being indirectly selected for. Thus, a comprehension of the longitudinal aspects of genetic (co)variances allows us to better understand the current status of novel traits from the time they start to be recorded, based on the past breeding decisions and current genetic correlations.

The expectation that selection can change genetic correlations over time implies that such correlations are longitudinally dynamic parameters. Thus, selection is not only changing the traits’ populational means, but also the genetic relationship between the traits. Thus, we hypothesized that changes in genetic correlations may represent changes in an underlying physiological trait responsible to regulate the expression of multiple traits [22]. While measuring genetic correlations as a populational parameter is indeed effective, the possibility to obtain individual-specific values of genetic correlations can further the potential for genetic gain in multiple traits in a breeding program.

In this work, the objectives were to **(1)** extend the inferences about how selection and random drift affect genetic variances [3] to how selection and random drift affect genetic correlations, by deriving the formulas for the changes in the (co)variance parameters due to multi-trait selection; **(2)** define an alternative method to estimate genetic correlations at the individual, rather than at the populational level, named by us as the *individualized sire genetic correlation*; and **(3)** verify if the theory from the first objective was in agreement with the empirical trajectory of genetic correlations observed over 30 years of data from the French Holstein dairy cattle population, comprised the trajectory of the individualized sire genetic correlation, while interpreting the trajectories with information from the breeding strategies deployed in the period that comprises the data.

## Methods

### Genetic evaluation model, assumptions, and the genetic correlation

We assume initially a two-trait framework, for the simplicity of the statistical derivations, and later describe how to extend the results to any number of traits. Let 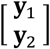 be the vector of phenotypic records for two traits, distinguished by the sub-indexes 1 and 2. Without loss of generality, we assume that all individuals have phenotypic records for both traits, and that phenotypes have been adequately pre-corrected for every non-genetic effect, being in the form of yield deviations. Thus, the two-trait animal model for genetic evaluation is given by,

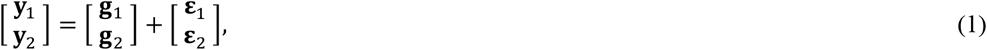

in which 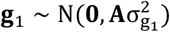 and 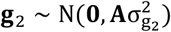 are the vectors of additive genetic effects (often referred to as breeding values in animal and plant studies), with 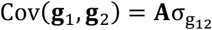, such that A is a relationship matrix between individuals (numerator, genomic, or combined pedigree and genomic [23, 24]); 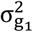 and 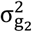 are the additive genetic variances, and 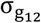 is the genetic covariance between the two traits; 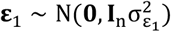 and 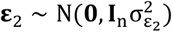 are the random residuals, with 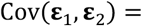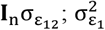 and 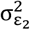 are the residual variances, and 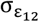 is the residual covariance. In this study, our interest is solely on the (co)variance parameters associated to the total additive genetic effects.

We now define 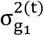 and 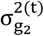 as the total additive genetic variances of trait one and two, respectively, at the t-th generation; 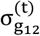 and ρ^(t)^ are respectively the genetic covariance and correlation between the two traits at the same t-th generation. Now, to simplify notation, let 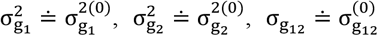, and ρ ≐ ρ^(0)^ be the initial variances, covariance, and correlation parameters at the base population (that is at the generation 0). It is widely known that genetic variances are affected by both selection and random drift [1–4], and it has been demonstrated that [3, 6],

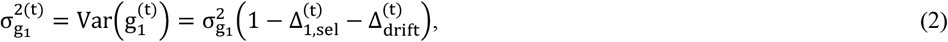

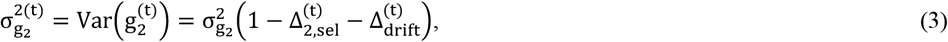

in which the terms Δ^(t)^ represent the cumulative changes in 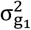 and 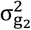 (the variances at the base population), over t generations, due to selection (sub-indexed sel) and random drift (sub-indexed drift).

Now, when A is the numerator relationship matrix, its diagonal entries are in the form A_ii_ = 1 + F_i_, in which F_i_ is the individual’s inbreeding coefficient. The expected change in genetic variances due to random drift in generation t is defined as 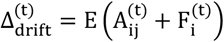, and can be directly estimated from the numerator relationship matrix [3]:

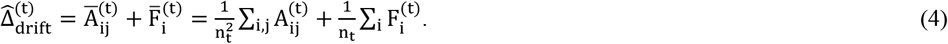

Therefore, if one estimates 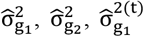 and 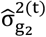, the changes in genetic variances due to selection in generation t are given by 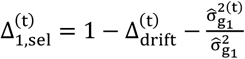 for the trait 1, and 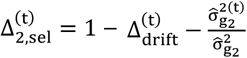 for the trait 2.

It is reasonable to assume that the genetic covariance is also affected by both selection and random drift. Thus, equations (1) and (2) can be extended to the genetic covariance as follows:

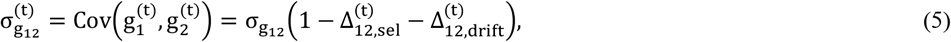

in which the terms 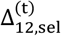 and 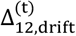 represents the cumulative changes in 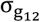.

Equations (2-3) and (5) are then combined to define the genetic correlation at the t-th generation,

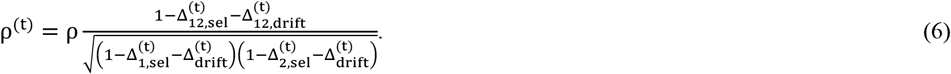

### Changes in genetic (co)variances due to selection on two traits

We move on now to explicitly defining the terms 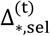 in equation (6), and we do so, based on the selection index (SI). When selecting individuals for two traits, the interest lies in maximizing their SI, which is defined at the t − th generation as:

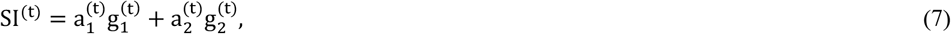

such that both 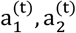 are weights defined by the breeding goals, economical indexes, and the genetic correlation between the two traits, considering the joint probability distribution of 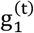 and 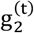. Although in principle, weights 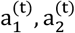 can assume negative values if the interest lies in selecting against undesirable traits (*e*.*g*. somatic cells or methane emission), we can assume 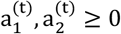 without loss of generality, since that would be equivalent to applying the SI on undesirable traits with their breeding values multiplied by minus one, and theoretical results are not at all changed.

Using the results from Appendix A and B, which present the moments of the truncated bivariate normal distribution [28], from equations (B8-B10), 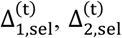, and 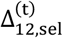 in equations (2-3) and (5) are given as:

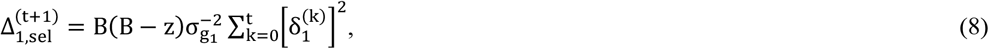

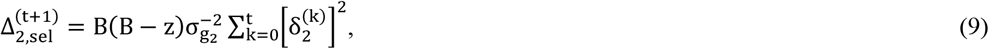

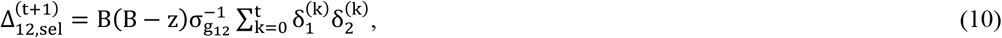

in which 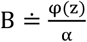, such that φ(z) is the density of Z ∼ N(0,1) at the point 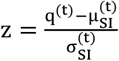, with the terms 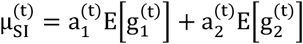 and 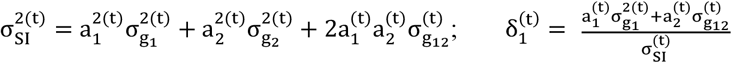 and 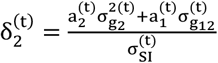 for any t. Since all 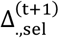 in equations (8-10) depend on summations for k = 0, …, t, in order to maintain these formulas valid for any t = 0,1,2, … we changed the super-indexes to t + 1, so that the summations for every 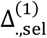 are valid. Although having 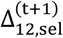 as defined in equation (10) dependant on 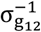 would require that 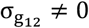 strictly, this is only an algebraic feature, so that 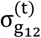 could be written as in equation (5), and we will show that 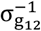 will be completely eliminated from ρ^(t+1)^ in equations (12-13), ensuring the validity of the theoretical derivation for any value of 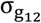, zero included.

### Changes in genetic covariance due to random drift

While changes in genetic variances due to random drift have been well described by [3], and presented here in section 2.1, the change in genetic covariance remains uncalculated explicitly. [3] affirm that the calculus path to obtain the changes in the genetic covariance due to random drift is analogous to that detailed in their work for the changes in the genetic variance. While the effect of random drift on the genetic covariance is linear, similarly to its effect on the genetic variances, as per equations (2-3), the final effect of random drift on the genetic correlation is non-linear, just as is the effect of selection, as per equation (6). This non-linear effect of both selection and random drift as drivers of changes in genetic correlations makes it harder to simultaneously derive both changes analytically. Thus, we opted to fully derive the changes in genetic correlation due to selection, *i*.*e*., a non-linear change explained as a function of 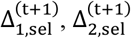, and 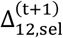, and consider any deviation from changes due to selection as changes due to random drift.

### Trajectory of the genetic correlation

We shall now generalize the trajectory of the genetic correlation between two traits in a population under selection, by replacing the terms 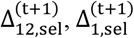, and 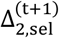 with their equivalences in equations (8-10), in equation (6), and we remind the reader that the super-indexes were changed to t + 1, so that the summations for every 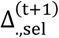 in equations (8-10) are valid. Thus, we have that:

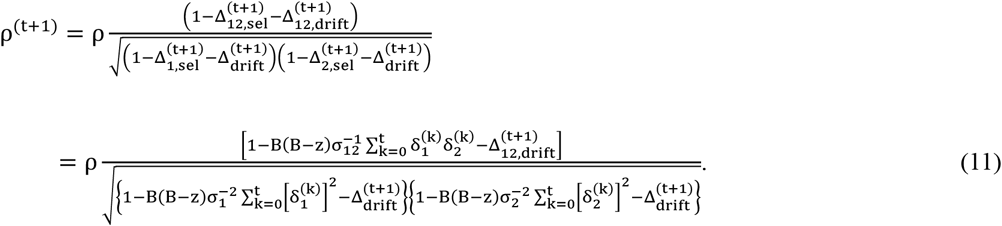

Now, as per equation (11), separating the changes in the genetic correlation due to selection and random drift is not straightforward, and we did not derive a closed form for 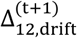. We will thus disregard any change due to random drift (*i*.*e*., we shall assume 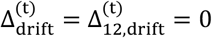, corresponding to 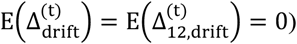. Moreover, to simplify the generalization of the mechanism of ρ^(t+1)^ for any t, we shall start by considering the change on a single generation:

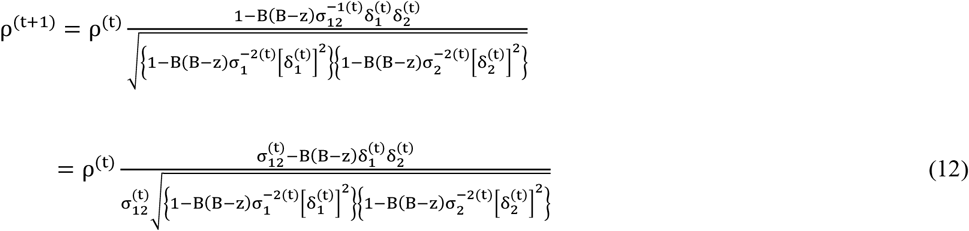

From the analysis of ρ^(t+1)^ as a function of ρ^(t)^, in Appendix C, we have that based on the selection index 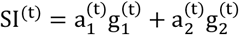, the new genetic correlation will be **(1)** ρ^(t+1)^ = ρ^(t)^, for 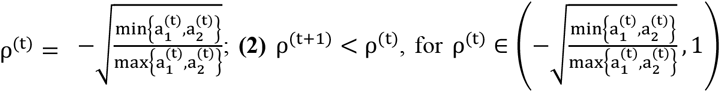 and **(3)** ρ^(t+1)^ > ρ^(t)^, for 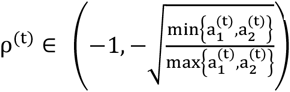. Therefore, in a recursive dependency, the value of ρ^(t+1)^ depends on 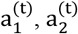 and ρ^(t)^, which in turn has its value depending on 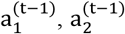 and ρ^(t−1)^, until we reach ρ^(1)^ that depends on the initial correlation 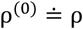 and on weights 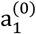 and 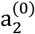. If we assume that 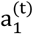 and 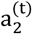 are fixed for all generations, *i*.*e*.,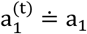 and 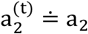, then, if:

**(1)** 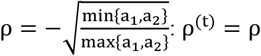 for every t.

**(2)** 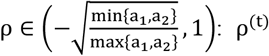 will decrease until a given generation t_1_, in which 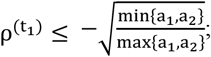

**(2.1)** 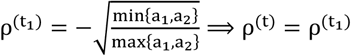for every t > t_1_;

**(2.2)** 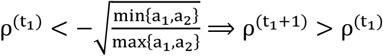, and the values of ρ^(t)^ will oscillate around, or converge to 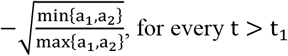.

**(3)** 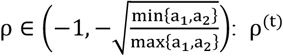 will increase until a given generation t_1_, in which 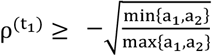

**(3.1)** 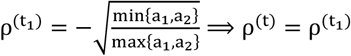 for every t > t_1_;

**(3.2)** 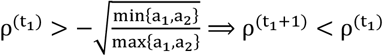, and the values of ρ^(t)^ will oscillate around, or converge to 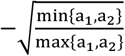, for every t > t_1_.

Before proceeding to the interpretation of the trajectory of ρ^(t)^ we must comment on the meaning of ρ decreasing or increasing, which varies according to the sign of ρ at the base population. For positive base population genetic correlations (ρ > 0), decreasing means that the correlation is attenuating, and increasing means that the correlation is intensifying. Now, for negative base population genetic correlations (ρ < 0), decreasing means that the correlation is intensifying, and increasing means that the correlation is attenuating.

The conclusions that can be drawn from the analysis above are that positive correlations will inevitable be attenuated, and depending on the weights a_1_ and a_2_, and selection intensity, positive genetic correlations may revert to negative; null genetic correlations will become negative; and negative genetic correlations have their trajectories determined by the weights a_1_ and a_2_, with potential to be either attenuated or intensified, and while in theory they do have the possibility to be become positive, the weights a_1_ and a_2_ must be carefully tailored for that purpose, and once reverted to positive, the natural path is to return towards zero and negative values.

Finally, we can account for the random drift as any deviation 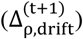 in the trajectory of the genetic correlation due to selection, and thus, we can re-write equation (12) as:

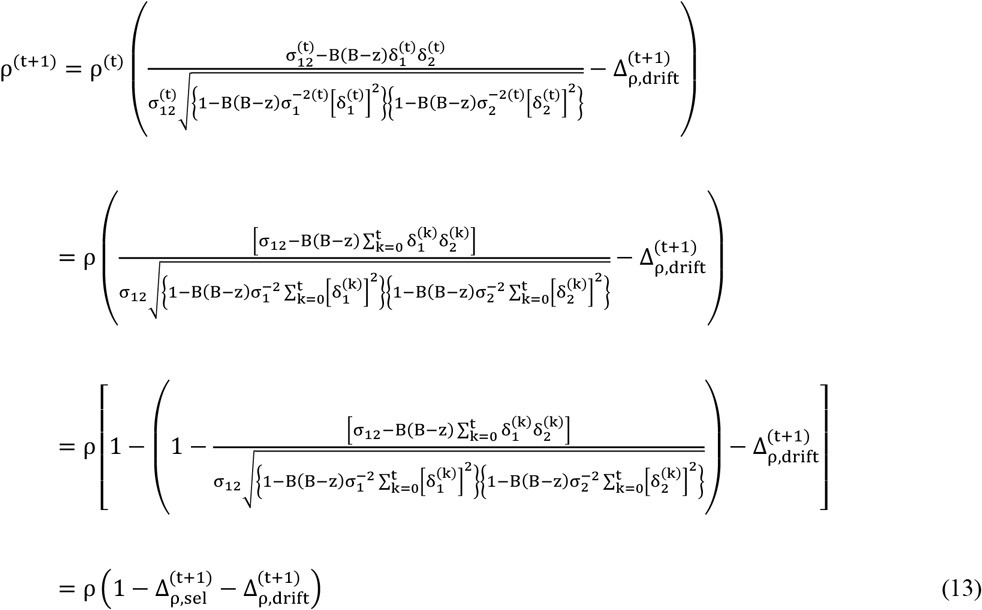

such that 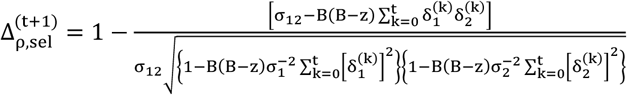. Note that if selection is not in place 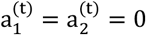, then 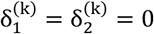 for every k = 0, …, t, and 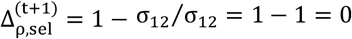, so that 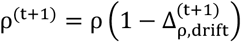, *i*.*e*. any deviation from the genetic correlation at base population, is purely due to random drift. It is important as well, to keep in mind that equations (11-13) require that genetic variance is never completely lost, *i*.*e*. that 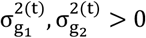, a requirement also imposed for the feasibility of selection in breeding populations.

### Changes in genetic correlation due to indirect selection and extension of theory to the selection on any number of traits

One must bear in mind that not only the direct selection for two traits will have an effect on the genetic correlation between them. Assume a population is under selection for two traits using the selection index 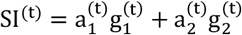 defined in equation (7). Any third trait not included in the SI will be indirectly selected for, when correlated to the traits directly selected for by the SI. Therefore, the genetic correlations between a third trait and the two traits included in the SI will also be impacted by the selection.

To detail the consequence of indirect selection to genetic correlations, we will assume three correlated traits. Now, the additive genetic effects of these three traits at generation t are 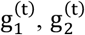 and 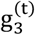, such that 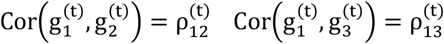, and 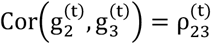. To understand how the selection for traits 1 and 2 affects the genetic correlation between traits 1 and 3, we redefine 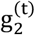 as the following linear combination of 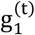 and a random variable Z_1_:

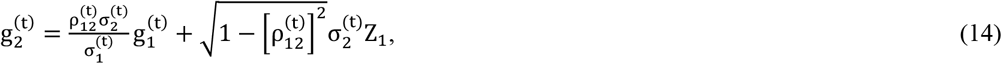

such that 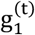 and Z ∼ N(0,1) are independent, ensuring that 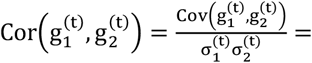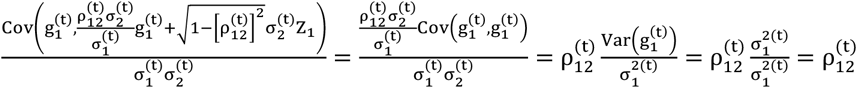. Now, since 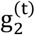 and 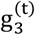 are correlated, it is necessary that 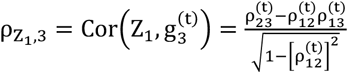 to satisfy 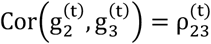. Thus, using the same structure from equation (14), it follows that 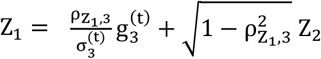, such that Z_2_∼ N(0,1) is independent from 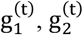 and 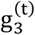, and:

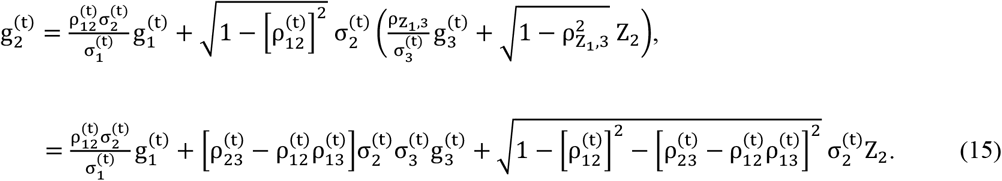

From equation (15), the SI can be redefined as:

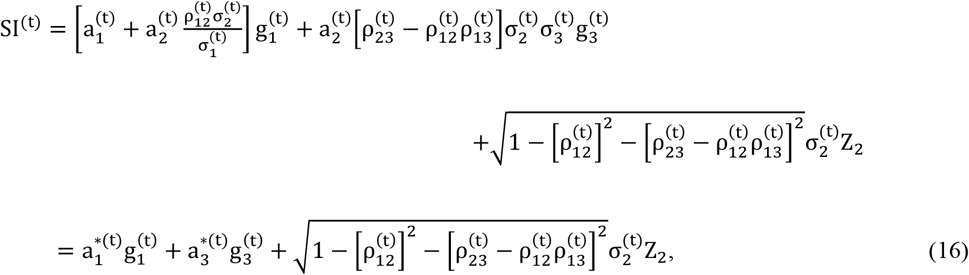

such that 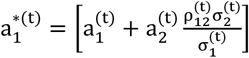, and 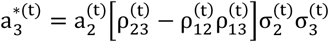 therefore, a selection using 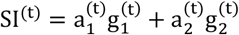 will impact the genetic correlation between traits one and three 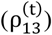 according with the rules defined in the subsection *Trajectory of genetic correlations*, depending on the value of 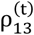 with respect to 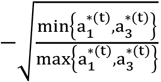. The impact on the genetic correlation between traits two and three 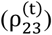 is analogous to that on 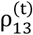, and can be understood by permutating sub-indexes 1 and 2 in 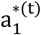 and 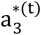 and equation (16).

Finally, we will generalize the effect of direct and indirect selection to any number of traits in a SI, since in real breeding programs the SI often involves more than two traits. This generalization allows us to extend the theory for the trajectory of genetic correlations to scenarios where selection is applied to any number of traits. Consider 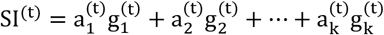, and using the structure from equation (15), we have that, for every j = 3, …, k:

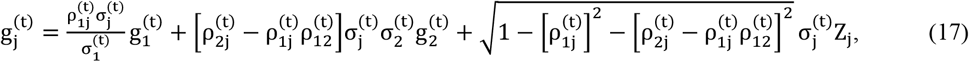

such that Z_j_ ∼ N(0,1) is independent from 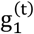 and 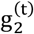. The SI can be then redefined as:

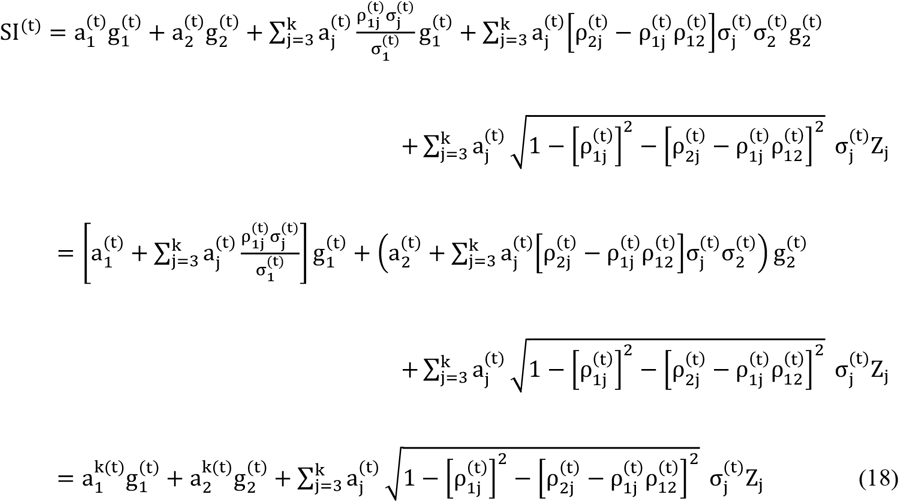

such that a 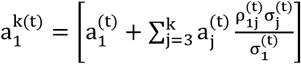 and 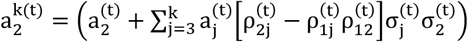, and therefore, a selection index 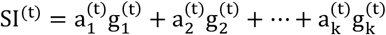 will impact the genetic correlation between the pair of traits one and two 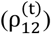 according with the rules defined in the subsection *Trajectory of genetic correlations*, depending on the value of 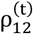 with respect to 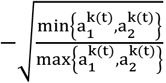.

Note that the result in equation (18) is general for any pair of traits in a multi-trait selection scheme. Moreover, even if neither trait 1 nor 2 are directly included in the SI, their genetic correlation will still be affected if each is correlated to at least one the traits included in the SI. The exemplify this, say 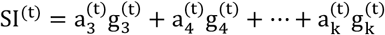. Therefore, the trajectory of the genetic correlation between traits 1 and 2 is ruled by 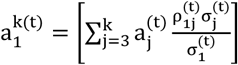 and 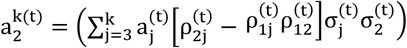, both different from zero if at least one 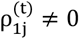, and one 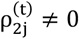.

### Real data set to study the trajectories of genetic correlations

A data set from the French Holstein dairy cattle population was used for our case-study. Our models were implemented on phenotypes in the form of yield deviations (YD), available for 4,501,624 cows born between 1991-2020, with a pedigree file containing a total of 8,275,018 animals that traced back three generations from the cows with performances. Figure 1 shows the number of cows with performance records per year of birth. All the cows in our models had YD available for all of the five following traits considered for the study: milk and protein yield (MY and PY), milking speed (MSPD), somatic cell score (SCS), and cow conception rate (CR, measured as success/failure on lactating cows, *i*.*e*. heifers excluded). The YD were output from the French national genetic evaluation, which evaluates MY, PY, SCS as the total performance per lactation, corrected for the duration; performance records comprise all lactations records per cow, and the model accounts for the repeatability (*i*.*e*., for the permanent environment of the cow). The five traits were chosen, so that we could study the trajectories of genetic correlations between traits with different signs of correlations (positive, negative, or almost zero), and between traits with different intensities of selection. For example, MY and PY are traits with a typically very high positive genetic correlation (∼0.9), both heavily selected for since the 1970’s. While SCS presents a very low genetic correlation with the production traits (∼0.03), it is a relevant trait, included in the French genetic evaluation and breeding goals for the Holstein dairy cattle since 1997 and 2001, respectively. Moreover, SCS presents a negative correlation with CR (∼-0.25). Both MY and PY are also well known to present a negative genetic correlation to CR (∼-0.2), with CR featuring in the French breeding goals for the Holstein dairy cattle since 2001. Finally, MSPD is a trait hardly selected for, with very low genetic correlation with MY, PY, and CR, but that presents a fair positive genetic correlation with SCS (∼0.35). The genetic correlations between the traits are approximate values made available by the French association GenEval (https://www.geneval.fr), responsible for performing the national genetic evaluation of the French Holstein dairy cattle population.

**Figure 1:**
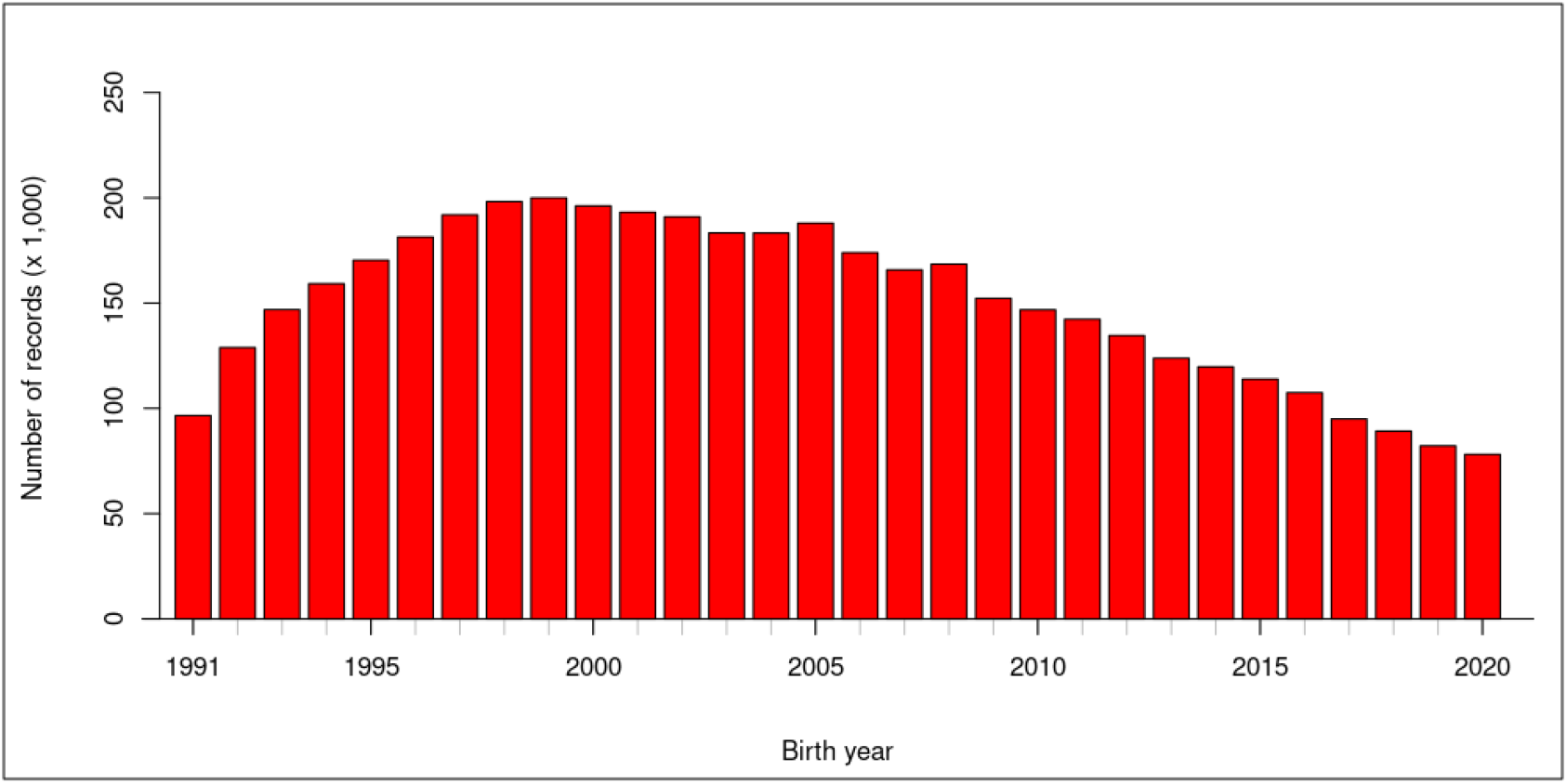
Number of cows with performance records available per year of birth, totalizing 4,501,624 cows.

### Genetic evaluation on the real data

The animal model was used to perform two-trait genetic evaluations for every pairwise combination of the five traits considered for our study. Since performances were in the form of YD, the model was as defined in equation (1), with the numerator (*i*.*e*. pedigree-based) relationship matrix defining the covariance structure A between the breeding values. Rather than using a frequentist approach for the genetic evaluation, *i*.*e*. the restricted maximum likelihood (REML) method [29, 30] to estimate variance components and Henderson’s mixed model equations [31, 32] to obtain the best linear unbiased predictors (BLUP) of the breeding values, we deployed a Gibbs sampler to estimate genetic parameters and predict breeding values, using GIBBS3F90 from the BLUPF90 family of programs [33]. This choice for the Gibbs sampler was based on the study from [6], who used a Gibbs sampler to estimate trajectories of genetic variances for 30+ years of data in the Manech Tête Rousse dairy sheep population. Therefore, we chose to deploy the same method to study the trajectory of genetic correlations between traits, and we detail in the next subsection how the estimates of genetic variances and correlations by year of birth were obtained from the Gibbs sampler. We verified the convergence of the Markov-Chain Monte-Carlo (MCMC) generated by the Gibbs sampler using the R [34] package Bayesian Output Analysis (boa) [35], using a graphical assessment assisted by Geweke’s criterion [36].

### Estimation of genetic variances and correlations by year of birth

The use of the Gibbs sampler allows to generate samples of genetic correlations per birth year of the animals, calculating the genetic correlations from the estimated breeding values (EBVs) at each of the MCMC samples kept. We used these estimates to observe the trajectories of genetic correlations between the selected traits in our population from the French Holstein dairy cattle. The method proposed by [6] was originally applied to a single-trait model, and consisted in:

1. Define the number of iterations (N), the burn-in (b), and the thinning (k) of the Gibbs sampler;
2. For every one of the j = 1, …, (N − b)/k MCMC samples kept, gather the 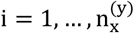 EBVs (ĝ_ij_) of either the males (x = M) or females (x = F) born in year y, for every birth year available, and compute the genetic variances as:

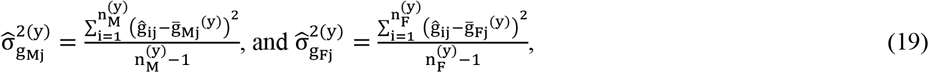

such that 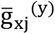 is the mean EBV of males/females born in year y;
3. Using the 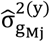 and 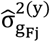 computed for the j = 1, …, (N − b)/k MCMC samples, obtain the final estimates of the genetic variances as:

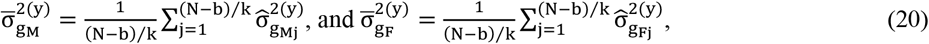

and their variances can be estimated as:

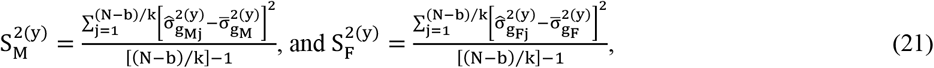

so that 95% confidence intervals can be drawn as:

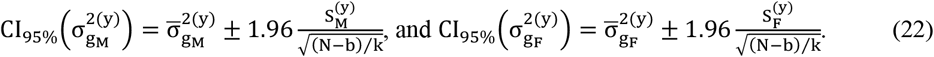

Confidence intervals for 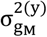 and 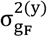 can also be drawn from the posterior distribution, directly from the quantiles 0.025 an 0.975. Nonetheless, the confidence intervals in equation (22) are valid for a sufficiently large (N − b)/k MCMC samples kept, since in this case the distributions of both 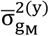 and 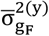 will converge to Gaussian.

We extended this method to compute genetic covariances in the two-trait models, enabling us to compute the genetic correlations. Therefore, for every one of the j = 1, …, (N − b)/k MCMC samples kept, we computed:

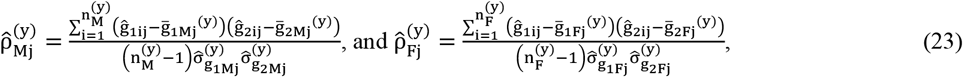

in which the sub-indexes 1 and 2 indicate the two different traits, and obtained the final estimates of the genetic correlations for each birth year y as:

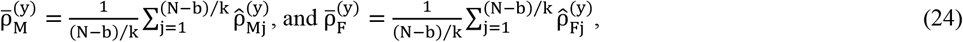

With posterior variances:

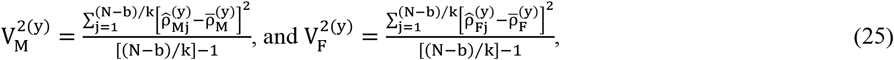

so that 95% confidence intervals could be drawn as:

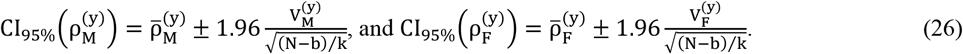

The following values of MCMC parameters were defined for the Gibbs sampler implemented to perform our case-study: we ran N = 300,000 iterations, discarding the burn-in of the first b = 100,000 iterations, and applied a thinning of k = 200 iterations, finally resulting in (N − b)/k = 1000 samples to compute the genetic correlations per year of birth, separately for the males (bulls or sires) and females (cows).

Given the large volume of data, in order to speed-up and ensure the convergence of the Gibbs sampler after the burn-in, we provided initial values for the variance components based on the heritabilities and genetic correlations used for the national genetic evaluation of the French Holstein dairy cattle.

Finally, results presented for the five traits individually were based on the combined results for each of their pairwise evaluation (each of these pairwise evaluations resulting in 1000 MCMC samples). Therefore, each trait was evaluated four times, and the individual trait results for their EBVs and heritabilities are presented based on the total 4000 MCMC samples from the four different Gibbs samplers that involved each trait.

### Individualized sire genetic correlation

Our last objective was to define a method to compute genetic correlations at the individual, rather than at the populational level. To do so, we defined what we called an individualized sire genetic correlation (iSGC) between pairs of traits. This consisted of a genetic correlation between traits, computed exclusively for sires, using the EBVs of their daughters corrected for the maternal EBV. The correction of the daughters’ EBVs for the maternal EBV was intended to ensure that the iSGC indeed captured the genetic background from the sires only. The choice to compute an individualized genetic correlation for sires only was because in dairy cattle populations, only sires have a large amount of offspring, so that an individualized genetic correlation can be assessed with reasonable accuracy. To obtain the iSGC as a genetic correlation expressed by the offspring, we proposed the following methodology:

1. Select sires with a sufficiently large number of daughters with phenotypic records;
2. For the i = 1, …, n_s_ daughters of each selected sire s, correct their EBVs for that of their dams, *i*.*e*.:

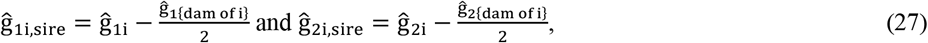

in which the sub-indexes 1 and 2 indicate the two different traits;
3. For each sire s, gather its i = 1, …, n_s_ daughters, and calculate the iSGC as:

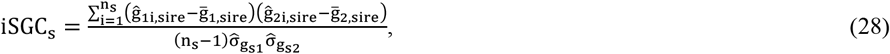

such that 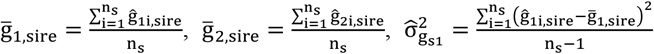, and 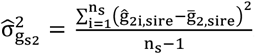.

The iSGC can be computed for each sire using either a REML-BLUP framework [29–32], from the single set of EBVs output from this method, or using the Gibbs sampler. Since we used the Gibbs sampler, similar to the method used to estimate 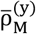 and 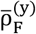 for bulls and cows per year of birth, we computed the iSGC for each one of the (N − b)/k = 1000 MCMC samples kept from our Gibbs sampler, and finally defined 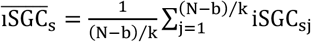.

With our data set, we evaluated sires born between 1991-2015, and for the calculus of the iSGC, we chose only those sires with ≥ 500 daughters that had YD records and dam information. Our selection totalized n_sires_ = 1161 sires for which an iSGC was calculated. Figure 2 presents **(a)** the distribution of the number of daughters from the 1161 selected sires, and **(b)** the number of selected sires born per year, between 1991-2015.

**Figure 2:**
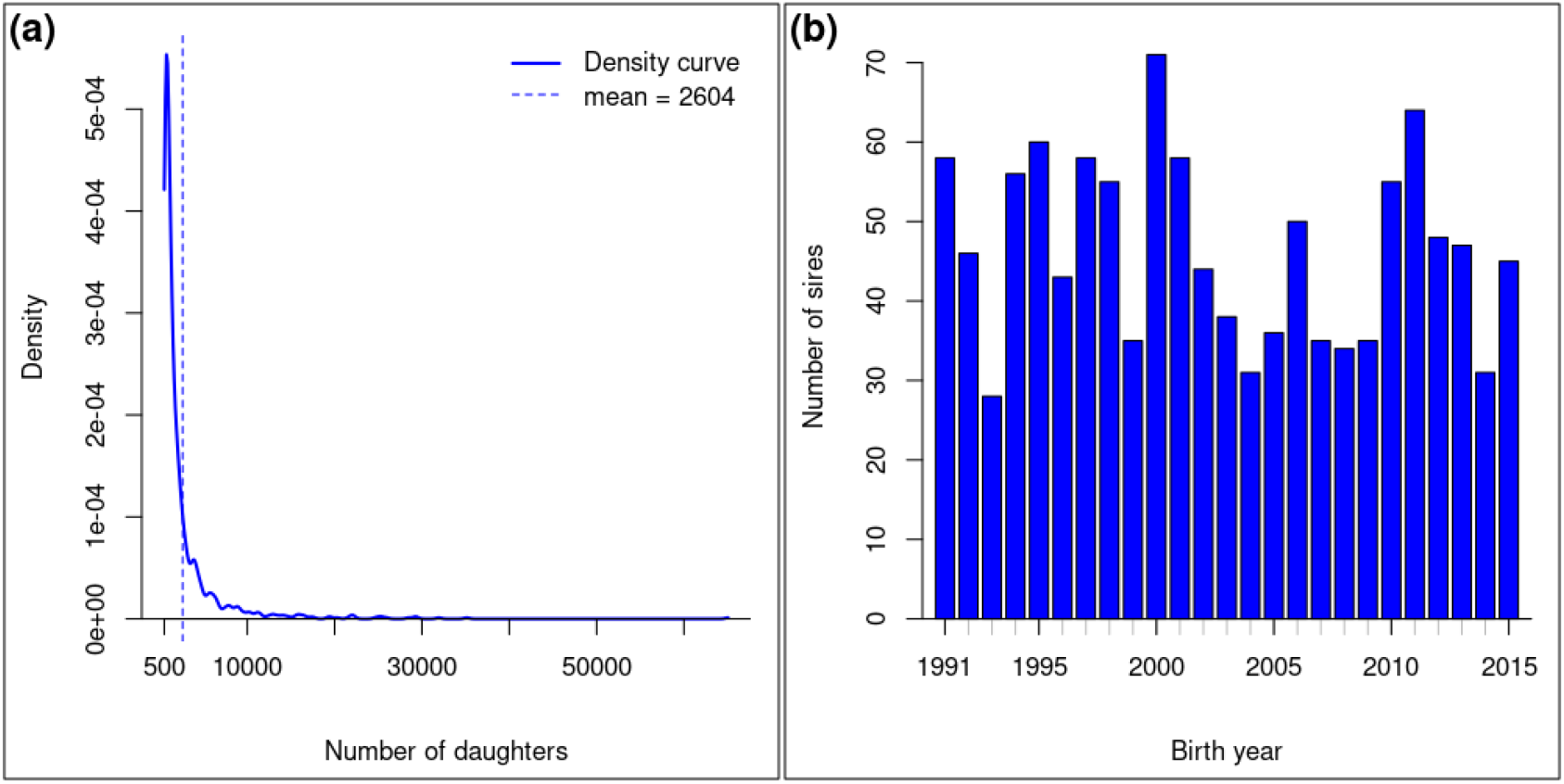
**(a)** Distribution (in the form of density function) of the number of daughters with performance records per selected sire used to compute the iSGC; **(b)** Number of selected sires born per year.

Since iSGC are unique for each sire, their populational summaries were computed as:

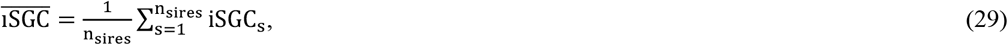

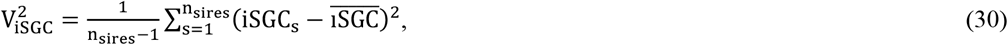

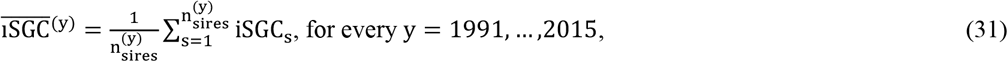

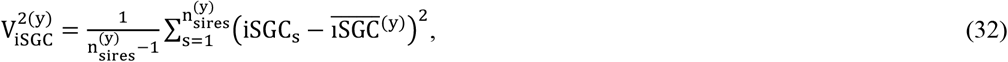

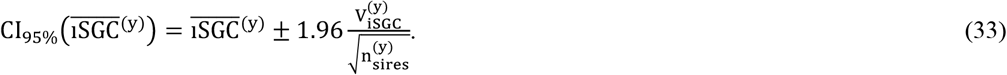

We compared the estimates of the genetic correlations obtained as parameters from the Gibbs sampler to 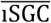, studied the distribution of all iSGC obtained for each of the 1161 sires selected for this analysis, studied the trajectory of the iSGC according to the birth year of the sires, using the summaries from equations (31-33), and compared the iSGC to the sires’ EBVs.

## Results

Since our study was separated in theoretical objectives and a case study, the results will be presented in the same framework. The theoretical results focus on the understanding and interpretation of the effect of selection on the trajectory of genetic correlations, and the case-study results present, among other summary statistics, the observed trajectories of genetic correlations between the pairs of traits studied, and the results related to the iSGC.

### Theory: the effect of selection on the trajectory of genetic correlations

From the theory derived in the materials and methods section, the genetic correlation between two-traits, after t + 1 generations under selection, can be expressed as 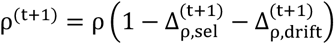, such that ρ is the original genetic correlation at generation zero,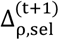is the effect of selection, and 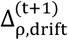 is the effect of the random drift.

The expression for 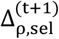 is fully detailed in equation (13), and the interpretation of these effects is that, disregarding random drift, in a population under selection, positive correlations will inevitably be attenuated (*i*.*e*. decrease), and depending on the weights a_1_ and a_2_ considered in the selection index 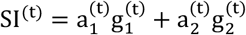 and on the selection intensity, positive genetic correlations may become negative; null genetic correlations will become negative; and negative genetic correlations have their trajectories determined by the weights a_1_ and a_2_, being either attenuated (*i*.*e*. increase) if their original values are lower than 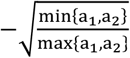, or intensified (*i*.*e*. decrease) if their original values are greater 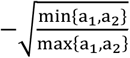. While in theory an originally negative genetic correlation may become positive, the weights a_1_ and a_2_ must be carefully tuned to achieve this result, and one must bear in mind though, that, once a positive value is achieved, the natural path of this genetic correlation will be to return towards negative values, revolving around 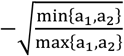.

These theoretical results, as well as their interpretation, are valid when selection is applied for any number of correlated traits, as detailed in the subsection *Indirect selection and changes in genetic (co)variances extended to selection on any number of traits*. To conclude the results on the theoretical aspect of our study, any change in the breeding goals (*i*.*e*., on the weights a_1_ and a_2_) will redirect the trajectory of the genetic correlation towards the new 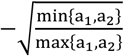, updated to the weights a_1_ and a_2_ according to the new breeding goals.

### Case-study

Table 1 presents a description of the five traits considered in our case-study, in the form of yield deviations (YD). Table 2 presents the heritabilities and genetic correlation estimates. Heritability estimates were computed using the Gibbs sampler variance components for each trait, by combining results from every two-trait analyses that considered the trait. A preliminary assessment (results not shown) showed no significant difference between the individual trait’s variance components and breeding values obtained with different traits combinations. The upper triangle of Table 2 presents the genetic correlation parameters from the Gibbs sampler on the entire population, while the lower triangle of Table 2 presents the mean 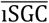 obtained for the 1161 sires selected to computed this parameter. Most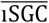 were not statistically different from the genetic correlations estimated on the entire population as a parameter from the Gibbs sampler. The two 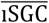 that differed statistically from their equivalent genetic correlation parameter from the Gibbs sampler were the pairs MSPD-SCS and SCS-CR, for which the Gibbs sampler parameters presented a more intense correlation than the 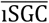.

**Table 1:** Description of the performance records available in the form of yield deviations, for the five traits evaluated.

| Trait | Description | Unit | Variance (unit<sup>2</sup>) |
| --- | --- | --- | --- |
| MY | Milk yield | Kg / lactation | 926,681.00 |
| PY | Protein yield | Kg / lactation | 954.00 |
| MSPD | Milking speed | Score: 1 (slow) to 5 (fast) | 0.60 |
| SCS | Somatic cell score | $\log_2(\text{SCC}/100,000) + 3$ | 0.69 |
| CR | Cow conception rate | % of success | 815.00 |

**Table 2:** Estimated genetic parameters (heritabilities and genetic correlations between pairs of traits), with their standard errors in parenthesis. Values in the diagonal (in bold) correspond to the heritability estimates; upper triangle values correspond to the genetic correlation parameters from the Gibbs sampler on the entire population; lower triangle values correspond to the mean iSGC 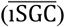 obtained for the 1161 sires selected to computed this parameter. 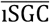 in bold are those that were statistically different from their equivalent genetic correlation parameter from the Gibbs sampler (significance level of 0.05).

|  | MY | PY | MSPD | SCS | CR |
| --- | --- | --- | --- | --- | --- |
| MY | **0.223 (0.001)** | 0.78 (0.05) | -0.06<sup>a</sup> (0.06) | -0.04<sup>a</sup> (0.06) | -0.15 (0.03) |
| PY | 0.87 (0.02) | **0.378 (0.001)** | -0.07<sup>a</sup> (0.09) | -0.01<sup>a</sup> (0.05) | -0.20 (0.04) |
| MSPD | -0.02<sup>a</sup> (0.04) | -0.02<sup>a</sup> (0.04) | **0.235 (0.001)** | 0.31 (0.08) | -0.04<sup>a</sup> (0.04) |
| SCS | -0.11 (0.05) | -0.12 (0.04) | 0.10<sup>b</sup> (0.04) | **0.126 (0.001)** | -0.26 (0.08) |
| CR | -0.07<sup>a</sup> (0.05) | -0.11 (0.05) | -0.01<sup>a</sup> (0.03) | -0.06<sup>ab</sup> (0.04) | **0.006 (0.002)** |

<sup>a</sup> estimated value not significantly different from zero (significance level of 0.05)
<sup>b</sup> $\overline{\text{iSGC}}$ values that were statistically different from their equivalent genetic correlation parameter from the Gibbs sampler (significance level of 0.05)

Figure 3 presents the genetic trend of all traits evaluated through the years 1991-2020 (diagonal panels), as well as the trajectories of the genetic correlations between the pairs of traits (lower triangle panels), and bivariate trajectory of the genetic progress for the pairs of traits (upper triangle panels), through these three decades.

**Figure 3:**
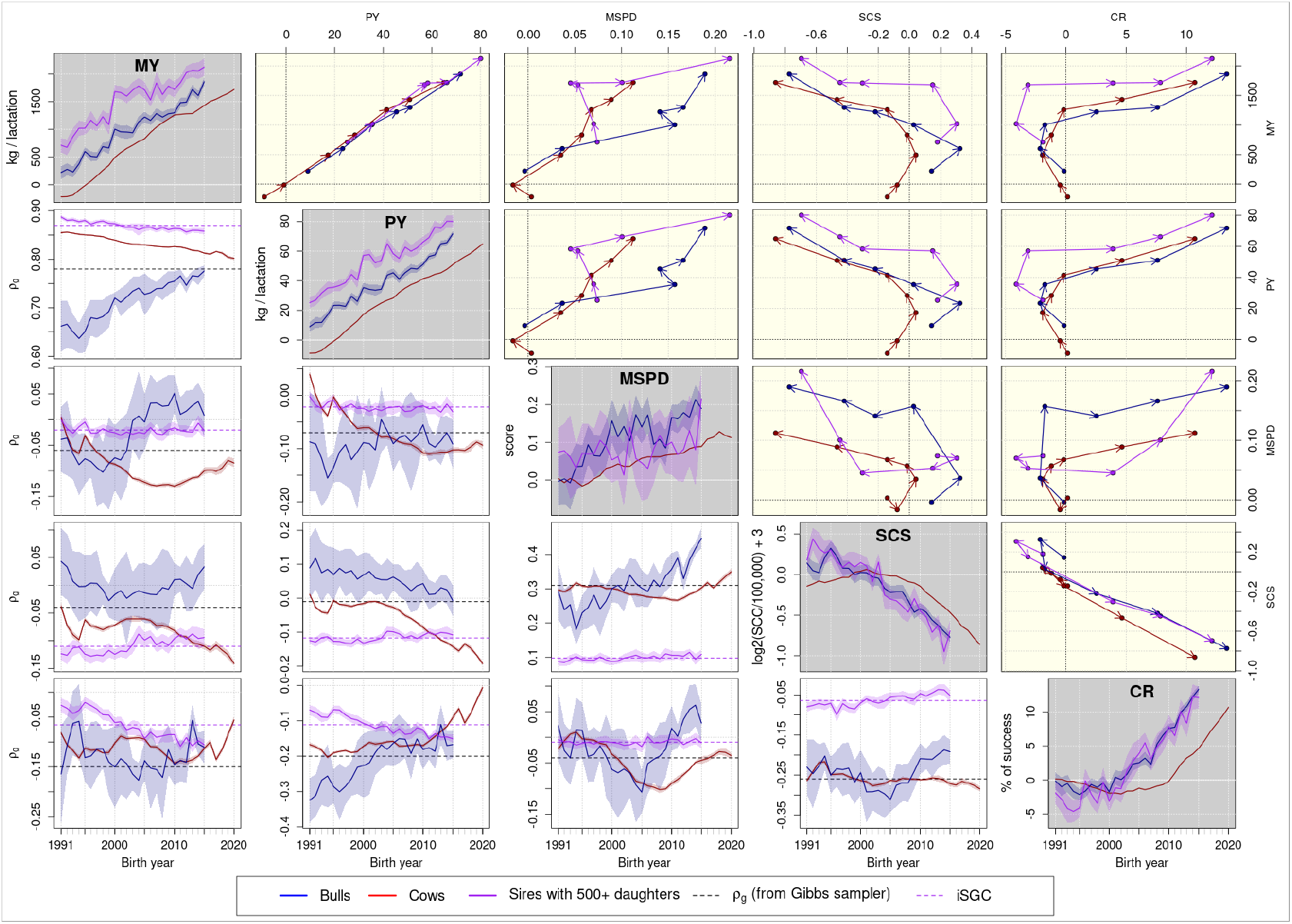
**(diagonal gray panels)** Genetic progress of all traits evaluated through the years 1991-2020, separated for bulls (all and only sires with more than 500 daughters) and cows, values are expressed as the differences from the population mean EBV at the year 1991; **(lower triangle white panels)** Trajectories of genetic correlations between the pairs of traits through the years 1991-2020, separated for bulls (all and only sires with more than 500 daughters) and cows; **(upper triangle light yellow panels)** Bivariate trajectory of the genetic progress for the pairs of traits evaluated through the years 1991-2020 for bulls (all and only sires with more than 500 daughters) and cows, with values expressed as the differences from the population mean EBV at the year 1991.

From the diagonal panels in Figure 3, we could observe that MY, PY, and MSPD showed a consistent increasing genetic trend, for males (blue and purple lines) and females (red lines), with the genetic trends from males being always higher than the trends from females. EBVs from the sires with more than 500 daughters (purple lines) were significantly greater than the average of all bulls for both MY and PY. The health trait (SCS) showed a consistent decreasing pattern for males (blue and purple lines), and a mildly increasing pattern for females (red line) born until 2001, inflecting from this year on and changing to a decreasing pattern until 2020. Until ∼2001, the EBVs from males were generally greater than the EBVs from females, however with the differences decreasing as the birth year approaches ∼1997-2004. After 2004, the EBVs from males became lower than the EBVs from females. The fertility trait (CR) showed a stable pattern for males (blue and purple lines), and a mildly decreasing pattern for females (red line), for animals born until 2001. From this year on, CR began to present a consistently increasing pattern for both males and females. Until ∼2001, the EBVs from males and females were in general not significantly different. After 2001, the EBVs from males became greater than the EBVs from females.

When analysing the trajectories of the Gibbs sampler parameters for the genetic correlations between the pairs of traits, presented in the lower triangle panels of Figure 3, the first thing noticed was that the trajectories were generally different for males (blue lines) and females (red lines). The exceptions for these differences were the trajectories for the genetic correlations between the pair MY-CR, for which both cows and bulls presented an overall flat pattern around ∼-0.12; between the pair PY-SCS, for which both cows and bulls presented a decreasing pattern running in parallel; and between the pair PY-CR, for which both cows and bulls presented an increasing pattern, although the figure suggests that from the year 2005, the genetic correlation between this pair of traits in bulls could be reaching an stable value. For the other pairs of traits, which presented different trajectories between males and females, bulls (blue lines) presented an overall increasing pattern for the pairs MY-PY and MSPD-SCS; flat pattern for the pairs MY-SCS and PY-MSPD; and an initially decreasing and then increasing pattern for the pairs MY-MSPD, MSPD-CR, and SCS-CR. Still with respect to the traits for which the trajectories of genetic correlations differed between males and females, cows (red lines) presented an overall decreasing pattern for the pairs MY-PY and PY-MSPD; flat pattern for the pairs MSPD-SCS and SCS-CR; an initially decreasing and then increasing pattern for the pairs MY-MSPD and MSPD-CR; and an initially stable and then decreasing pattern for the pair MY-SCS. Moreover, the trajectories of the Gibbs sampler correlation parameters (blue and red lines) differed substantially from the trajectories of the iSGC (purple lines), calculated for the 1161 bulls with more than 500 daughters. The iSGC presented an overall increasing pattern for the pairs MY-SCS, PY-SCS, and SCS-CR; decreasing pattern for the pairs MY-PY, MY-CR, and PY-CR; and flat pattern for the pairs MY-MSPD, PY-MSPD, MSPD-SCS, and MSPD-CR.

The bivariate trajectories of the genetic progress for the pairs of traits evaluated through the years 1991-2020, presented in the upper triangle panels of Figure 3, showed that as expected, breeding goals through this period have aimed to jointly select favorably most traits evaluated in our study.

Figure 4 presents the histograms of the iSGC obtained for the 1161 bulls with more than 500 daughters (lower triangle panels), and scatterplots of the EBVs obtained for these bulls, for every pair of traits evaluated (upper triangle panels). The histograms presented in the lower triangle panels of Figure 4 showed reasonable dispersion of iSGC around their mean value. The density lines drawn over the histograms for the years 1991, 1995, 2000, 2005, 2010, and 2015 reinforce the trends in the trajectories of iSGC already described by Figure 3, *i*.*e*., an overall increasing pattern for the pairs MY-SCS, PY-SCS, and SCS-CR; decreasing pattern for the pairs MY-PY, MY-CR, and PY-CR; and stable pattern for the pairs MY-MSPD, PY-MSPD, MSPD-SCS, and MSPD-CR, yet for these last four pairs of traits, the iSGC presents an increase in its dispersion in the more recent years. The scatterplots of the EBVs for every pair of traits evaluated (upper triangle panels), along with the 95% confidence ellipses for the years 1991, 1995, 2000, 2005, 2010, and 2015, confirm that the breeding goals through this period have aimed to jointly select favorably for all the traits evaluated in our study. In this figure, the 95% confidence ellipses aimed to assist the visualization of the subtle change in the genetic correlations between traits over the years.

**Figure 4:**
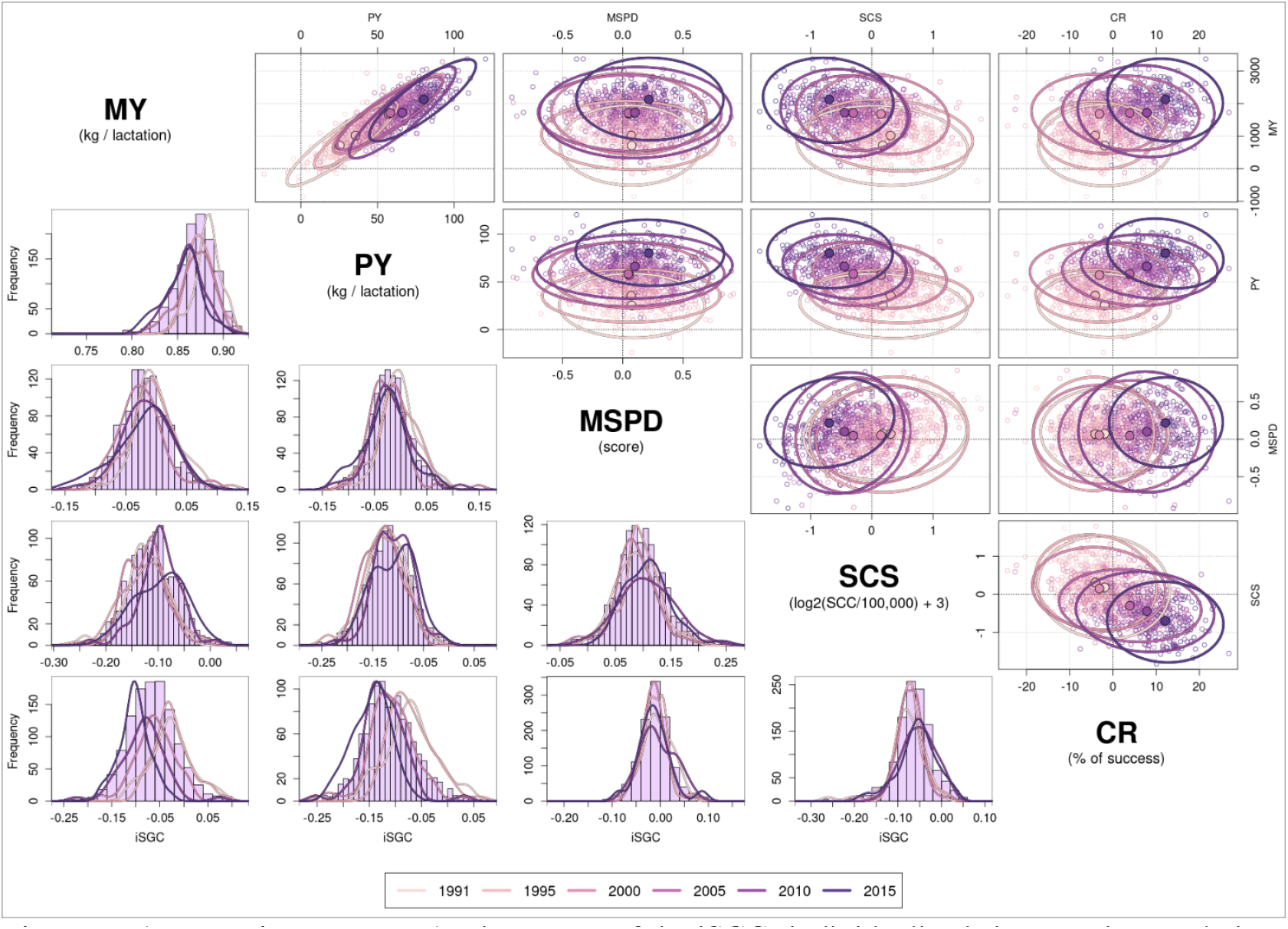
**(lower triangle panels)** Histograms of the iSGC (individualized sire genetic correlation) obtained for the 1,161 bulls with more than 500 daughters for all the pairs of traits evaluated, along with the densities calculated for the iSGC of bulls born in the years 1991, 1995, 2000, 2005, 2010, and 2015; **(upper triangle panels)** Scatterplots of the EBVs obtained for the 1,161 bulls with more than 500 daughters, for every pair of traits evaluated, along with the 95% confidence ellipses calculated for the EBVs of bulls born in the years 1991, 1995, 2000, 2005, 2010, and 2015, with values expressed as the differences from the population mean EBV at the year 1991.

Figure 5 presents the scatterplots of the EBVs obtained for the 1161 bulls with more than 500 daughters vs. their iSGC. Lower triangle panels present the scatterplots of the iSGC (y-axis) and the EBVs (x-axis) of the traits named above the panels, and upper triangle panels present the scatterplots of the iSGC (y-axis) and the EBVs (x-axis) of the traits named on the left of the panels. In Figure 5 we observed that while selection did result in a change of the iSGC for some pairs of traits, not always the iSGC are themselves correlated to the sires’ EBVs for the traits in question. We assessed the correlations between the iSGC and the EBVs by judging that a potential correlation existed when the correlation calculated on values corrected by the yearly means (*i*.*e*., centering all data in zero, however accounting for the yearly differences) were out of the (−0.1,0.1) interval. This assessment suggested that MY may be positively correlated to iSGC_MY,SCS_, and negatively correlated to iSGC_MY,PY_ and iSGC_MY,CR_; PY may be positively correlated to iSGC_PY,SCS_, and negatively correlated to iSGC_MY,PY_ and iSGC_PY,CR_; MSPD may be positively correlated to iSGC_MSPD,SCS_, and negatively correlated to iSGC_MY,MSPD_ and iSGC_PY,MSPD_; SCS may be positively correlated to iSGC_MSPD,SCS_; and CR positively correlated to iSGC_MY,CR_ and iSGC_PY,CR_, and negatively correlated to iSGC_SCS,CR_.

**Figure 5:**
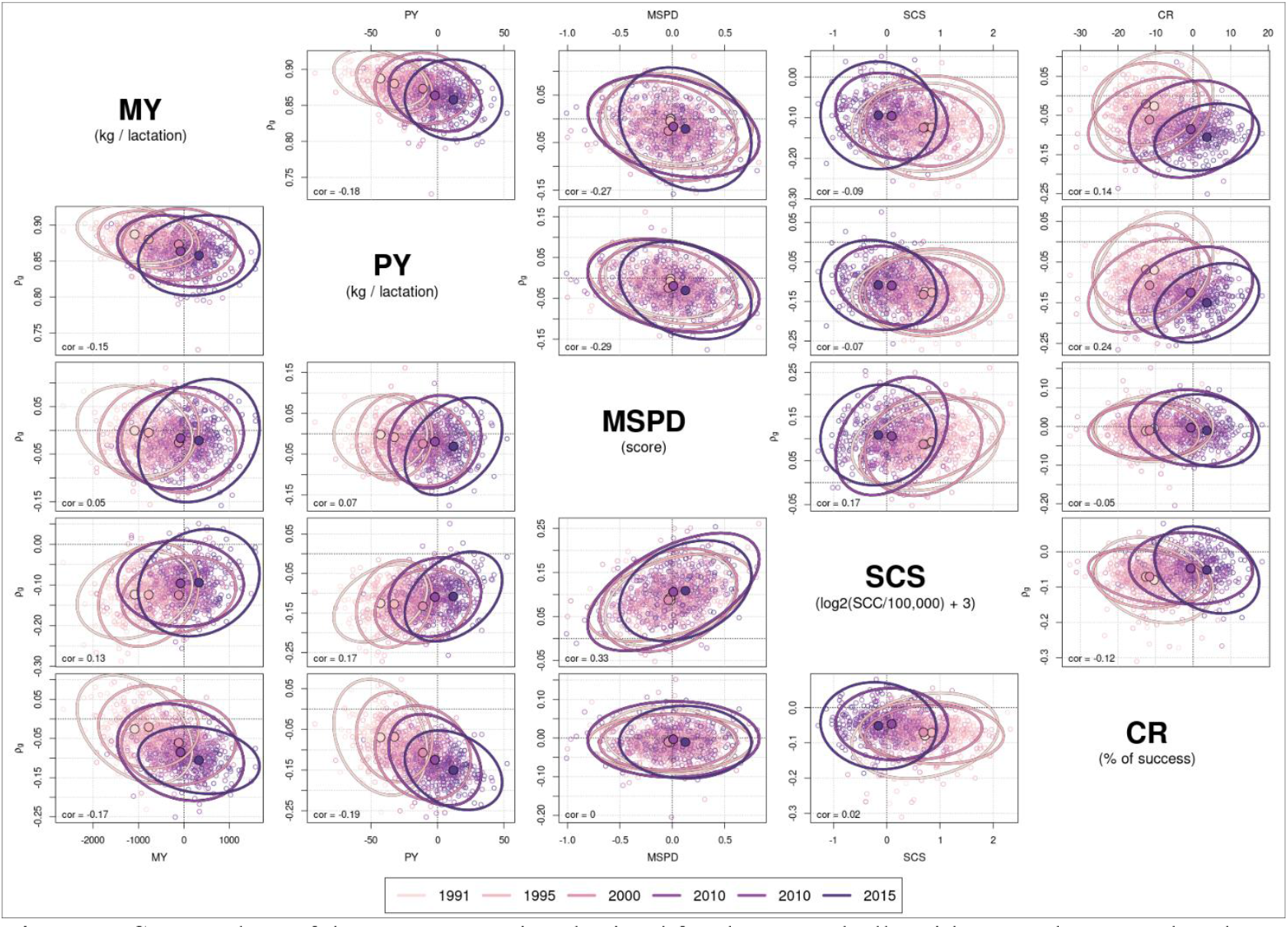
Scatterplots of the EBVs (x-axis) obtained for the 1,161 bulls with more than 500 daughters vs. their iSGC (individualized sire genetic correlation) (y-axis), along with the 95% confidence ellipses calculated for the EBVs of bulls born in the years 1991, 1995, 2000, 2005, 2010, and 2015.

## Discussion

Similar to the *Results* section, the discussion is presented in separately for the theoretical results, and the case study. However, different from the results, our theoretical discussion focuses on both the understanding and interpretation of the effect of selection on the trajectory of genetic correlations, and on the properties and meaning of the iSGC. The case-study discussion focuses on the most relevant results observed for the French Holstein dairy cattle population, relating them to the derived theory and to historical breeding goals in this population.

### Theory: trajectories of genetic correlations in populations under selection and iSGC

This study aimed, as a first objective, to understand how selection and random drift affect genetic correlations, by deriving the complete formulas for the changes in the (co)variance parameters due to a multi-trait selection scheme. Still on a theoretical framework, we had as another objective, which was to define a method to calculate genetic correlations at the individual, rather than at the populational, level.

We demonstrated that when a selection index 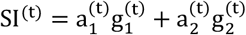 is applied on two traits, if this index is maintained throughout generations, the genetic correlation between the two traits involved will converge around 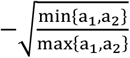, and the direction of its trajectory (increasing or decreasing) is determined by the original correlation. If a correlation is originally greater than 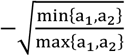, it will decrease towards this value, whereas if a correlation is originally lower than 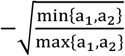, it will increase towards this value. Matching our results, [10] concluded that, if the SI is built with equal weights (*i*.*e*.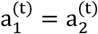), the original genetic correlation tends to a decreased value, and if selection is performed on a single trait only (*i*.*e*.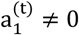 and 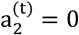 or vice-versa), the original genetic correlation will converge to zero. These conclusions from [10] match with ours, from the theoretical results presented in Figure C1, on the central panel (in which 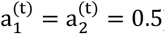) and on the top/bottom panels (in which 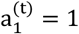 and 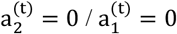 and 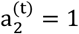), respectively.

Besides [10], other researchers have raised the question about the effect of selection on genetic correlations [3, 8, 9]. Most of these studies, however, focused on the direct consequences to genetic correlations due to changes in allele frequencies, and assumed a simpler model with very few loci as causative mutations. Our study shows that those results obtained under simpler genetic models may not be generalized to complex traits, which involve a very large number of causative mutations, either acting in pleiotropy, or with strong LD between causative mutations with an effect in the different traits. While the study from [10] is applicable to complex traits, its scenarios were limited to those aforementioned, assuming either equal weights for both traits involved in the SI, or selection for a single trait only.

Up to our knowledge, our study is the first to describe the changes in genetic correlations between traits under selection (direct or indirect) generalized for any values of weights in the SI, as well as accounting for multiple traits under selection. Inferences on the trajectory of genetic correlations can be highly relevant for breeding programs, since SI rely strongly on the genetic correlations. While non-updated populational correlations do not incur on misleading ranking of the selection candidates for a single trait, the same may not be true for the ranking of these candidates based on their SI, since individuals born in different years may present different genetic correlations regulating their genetic potential for two traits. Furthermore, given a SI, our theory allows the projection of future genetic correlations. Such projection may allow decisions on which selection candidates to choose, or even on how to redefine the SI, in order to avoid detrimental changes in genetic correlations between traits of commercial interest.

We have also proposed an alternative method to calculate genetic correlations between traits, that allows to obtain genetic correlations at the individual level, rather than the typically estimated genetic correlations as a population parameter. This proposal to calculate genetic correlations at the individual level was initially contemplated to address the changing genetic correlations over time. Another assumption, however, underlies in the fact that genetic correlations are not static parameters over time, in breeding populations. A change in genetic correlations due to selection means a change in the individuals’ physiological ability to regulate two or more traits altogether [37]. Thus, changes in genetic correlations could represent changes in an underlying physiological trait, which in turn may be thought of as a latent phenotype regulating multiple traits [22]. Although such latent physiological trait may be unmeasurable in practice, if genetic correlations can be measured at the individual level, they might work as proxies for such trait.

The approach to calculate genetic correlations at the sire level has been previously suggested in a study of the trade-off between production and fertility traits in a French Montbéliarde dairy cattle population [22]. This study, performed in a much smaller population compared to that of our current study, obtained genetic correlations for 247 sires with more than 500 daughters, similar to how we propose here, except that the daughters’ EBVs were not corrected for their maternal additive genetic effect. Nonetheless, the results on these 247 already suggested that different sires expressed different genetic correlations through their daughters, thus raising the discussion whether genetic correlations are indeed parameters, or a latent phenotype [22]. Although in the longitudinal perspective, the results from [22] suggest a negative genetic correlation mildly attenuating, opposite to our results obtained on a French Holstein dairy cattle population, the number of sires born per year in [22] is limited, and the longitudinal results must be evaluated with reservations, particularly given that the daughters’ EBVs were not corrected for their maternal additive genetic effect, which could blur longitudinal inferences.

We recognize that the calculus of iSCG is limited to populations in which individuals have a large number of offspring, such as breeding systems that focus on the productive females, *i*.*e*., systems for which selection is mostly performed on males, while production is achieved through the females, such as dairy cattle or dairy sheep. Nonetheless, whenever possible to calculate (*i*.*e*., whenever individuals have a sufficiently large number of offspring on which performances are recorded), the iSGC appears to be a trustworthy measure for genetic correlations, and a better method to evaluate their trajectories over time.

Once the iSGC is measurable, a rather wide dispersion on the distribution of the iSGC is a result with potential to suggest that genetic correlations may indeed be a non-static population parameter, identical for all individuals concerned in a genetic evaluation. Moreover, knowing that genetic correlations may be individual-specific values, given their nature of determining the relationships between traits, the iSGC may be thought of as the potential expression of an underlying trait responsible to regulate the observable phenotypes. This concept can be relevant when in the context of stabilizing selection [38], *i*.*e*., when the boundary of genetic progress is achieved by selection in a breeding program, thus limiting future potential gain. Accounting for individualized genetic correlations in a SI may provide an extra potential for further selection and genetic progress, by also allowing the selection of individuals better able to simultaneously regulate the traits of interest.

### Case-study

Our study aimed to connect the theory to a case-study in a real data set from the French Holstein dairy cattle population, comprising thirty years of phenotypic records, by interpreting the trajectories of genetic correlations with information from the breeding strategies deployed throughout the period of the records available in our data. We will focus our discussion about the results from the case-study on genetic correlations that had at least one of their estimates presented in Table 2 significantly different from zero, *i*.*e*., between the pairs MY-PY, MY-SCS, PY-SCS, MY-CR, PY-CR, MSPD-SCS, and SCS-CR.

Before further discussing properties of the iSGC, and the trajectories observed for the genetic correlations, it is relevant to make a remark about the EBVs observed for the 1161 bulls with more than 500 daughters, for which the iSGC were calculated. Particularly for the two production traits (MY and PY), the EBVs from the sires with more than 500 daughters were significantly greater than the average of all bulls. That is likely because these 1161 bulls are the elite bulls, and thus, the most used for inseminations, having the largest number of daughters.

In a preliminary study, iSGC were calculated directly from EBVs output from a BLUP genetic evaluation, that used REML estimates for the (co)variance components. Interestingly, iSGC based on the BLUP outputs were not very different from the iSGC based on the Gibbs sampler, and the correlation between iSGC_BLUP_ and iSGC_GIBBS_ was of 0.96. This result suggests that iSGC can be calculated directly as a sub-product from a routine genetic evaluation (which rarely uses Bayesian approaches, due to the computational burden demanded by such methods), without further great computational effort.

Overall, the mean iSGC were not significantly different from the population parameter estimates obtained by the Gibbs sampler. The exceptions were for the pairs of traits MSPD-SCS and SCS-CR, for which the values of the iSCG were less intense than the values from the Gibbs sampler, and we have not been able to objectively explain why such a difference was observed. Figure 4 presents the histograms of iSGC, and while genetic correlations are always treated as static parameters common to all individuals in a genetic evaluation, these histograms suggest that genetic correlations differ substantially between individuals, supporting our assumption that the iSGC can be considered as proxies of an underlying trait responsible to regulate the observable phenotypes measured in a production system.

In the lower triangle panels of Figure 3, we observed that the trajectories for the males differed substantially from the Gibbs sampler correlation parameters (blue lines) to the iSGC (purple lines), even though mean values of the iSGC were not significantly different from the population parameter estimates obtained by the Gibbs sampler. The iSGC was calculated as a sire-specific genetic correlation, expressed through the EBVs of their daughters. The correction of the daughters’ EBVs for the maternal breeding value, before correlating their breeding value to obtain the iSGC, aimed to ensure that the iSGC indeed captured the genetic background from the sires only. Moreover, differently from the classic genetic correlation parameter, which is estimated based on the average linear relationships (or lack thereof) between the breeding values of two traits, while accounting for the population structure, the iSGC searches for individual-specific patterns of correlations, and when measured on sires through the EBVs of their daughters, it also considers the recombination of the genetic background, being the iSGC a potentially less biased measure of genetic correlations between traits. Finally, the trajectories of the iSGC were in more agreement with the theoretical results than the genetic correlations inferred as a population parameter, and their confidence intervals were much narrower than those obtained for the trajectories inferred for the males from the Gibbs sampler correlation parameters. Considering all these properties of the iSGC, we considered them a better measure to study the trajectories of genetic correlations for the sires in our population, and all following discussions will be based on the purple lines of Figure 3, which represent the trajectories of the iSGC.

Overall, for the genetic correlations between the pairs of traits that had at least one of their estimates presented in Table 2 significantly different from zero (MY-PY, MY-SCS, PY-SCS, MY-CR, PY-CR, MSPD-SCS, and SCS-CR), we observed that sires (purple lines) and cows (red lines) presented different trajectory patterns, as per the lower triangle panels in Figure 3. The exception of these different trajectory patterns between the two genders was for the pair of traits MY-PY, for which both males (purple line) and females (red line) present a parallel decreasing pattern, as per the theory derived, with the values of the iSGC greater than those for the cows. While different trajectory patterns between the genetic correlations from the two genders may suggest different results in their evolution, it may also suggest differences from the estimation method. Given the properties from the iSGC discussed in the previous paragraph, we will focus interpretation on the trajectories of this correlation measure, which we deemed potentially less biased.

The genetic correlations between the pairs of traits MY-SCS and PY-SCS presented an initially stable and then decreasing pattern for the cows (red lines), and a rather stable pattern in their trajectories for the sires (purple lines), although a shift that mildly attenuated (*i*.*e*. increased) these negative genetic correlations was observed between 2000-2005, a likely consequence of the inclusion of SCS in the French breeding goals for this breed just before 2000.

While cows (red lines) presented an oscillating pattern around a mean of approximately -0.12 for the trajectory of the genetic correlation between the pair of traits MY-CR, and an attenuating (*i*.*e*. increasing) pattern for the pair PY-CR, approaching zero, sires (purple lines) presented a consistently intensifying (*i*.*e*. decreasing) pattern in their trajectory for these negatively correlated pairs of traits. These trajectories of the iSGC are in agreement with the theoretical results when one trait has a much greater weight in the SI than the other (the case for the French Holstein, as for many other dairy cattle populations, for which the weight on the production traits is much greater than on all other traits involved), and the initial genetic correlation is not more negative than the ratio 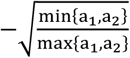, in which a_1_ and a_2_ are the weights attributed to the traits in the SI.

The genetic correlation between the pair of traits MSPD-SCS presented a generally flat, yet mildly increasing pattern for the sires (purple lines), and an overall flat pattern for the cows (red lines), although the pattern may be increasing after the year 2010. Both these traits have a rather low weight in the SI, compared to other commercial traits involved in the French breeding goal for the Holstein dairy cattle, and particularly MSPD is often not selected for by many producers. Along with a generally low genetic correlations between these two traits and the others included in the selection index, the dual selection for MDPS-SCS is not very intense, and can explain the rather stable pattern observed for the trajectory of the genetic correlation between these traits.

Lastly, the genetic correlation between the pair SCS-CR presented an overall flat pattern for the cows (red lines), and mildly attenuating (*i*.*e*. increasing) pattern for the sires (purple lines). Similar to the pair of traits MSPD-SCS, both SCS and CR had a rather low weight in the SI until 2012, compared to other commercial traits, being their selection not very intense on the period comprised by most of the data, thus explaining only minor changes in the trajectory of genetic correlations between the pair SCS-CR.

Interestingly, most of the iSGC present a correlation with the EBVs of their traits concerned, as shown in Figure 5, and we shall discuss this for every pair of traits that had at least one of their estimates presented in Table 2 significantly different from zero (MY-PY, MY-SCS, PY-SCS, MY-CR, PY-CR, MSPD-SCS, and SCS-CR), judging these correlations relevant when the ISGC and EBV presented a correlation out of the (−0.1,0.1) interval.

When it comes to the production traits (MY and PY), for which the mean iSGC was 0.87, both MY and PY were negatively correlated to iSGC_MY,PY_, suggesting that individuals with greater genetic potential for either one of these production traits are less able to maintain both production traits on high standards (thus, the dilution of proteins in milk). In other words, animals with the highest genetic potential for MY will compromise the PY, and animals with the highest genetic potential for PY will present a more limited genetic potential for MY. Just as selection tends to attenuate the iSGC (with its means values decreasing from 1991 to 2015), the 95% confidence ellipses in Figure 5 suggest that selection also attenuates the negative correlations observed between MY-iSGC_MY,PY_ and between PY-iSGC_MY,PY_, making both production traits less dependent on the genetic correlation that rules their relationship.

With respect to the relationships between the production traits and the health trait (SCS), for which the population genetic correlations between the pairs MY-SCS and PY-SCS were statistically zero, SCS was neither correlated to iSGC_MY,SCS_ nor to iSGC_PY,SCS_. However, MY and PY were positively correlated to iSGC_MY,SCS_ and iSGC_PY,SCS_ respectively, suggesting that an increase in the genetic potential for production does indeed lead to an increase in the genetic susceptibility to mastitis. From the 95% confidence ellipses in Figure 5, this relationship between MY-iSGC_MY,SCS_ and between PY-iSGC_PY,SCS_ seems to be maintained stable throughout the years.

Concerning the relationships between the production traits and the fertility trait (CR), for which the mean iSGC were -0.07 (statistically equal to zero) and -0.11 respectively for the pairs of traits MY-CR and PY-CR, MY and PY were negatively correlated to iSGC_MY,CR_ and iSGC_PY,CR_ respectively, while CR was positively correlated to both iSGC_MY,CR_ and iSGC_PY,CR_. The negative correlations between the production traits and the iSGC confirm that individuals with greater genetic potential for production are less able to balance the trade-off with fertility. One would expect that the same conclusion should be drawn for CR. However, the positive correlation between CR and the iSGC’s suggest that individuals with a greater genetic potential for fertility are also those better able to balance the trade-off with production, which may seem to contradict the statement that individuals with greater genetic potential for production are less able to balance the trade-off with fertility, and therefore we need to discuss these results carefully. When observing the 95% confidence ellipses in Figure 5, we can see that the positive correlation between CR and the iSGC’s is not very clear for bulls born before the year 2000, and that the negative correlation between the production traits and the iSGC is attenuating for bulls born after the year 2000. Fertility was included in the breeding goals for the French Holstein dairy cattle in the year 2001, which means that the bulls selected to match with the updated breeding goals were generally born from the year 2000 on. As selection considered both production and fertility, one could say that indirectly, individuals were selected to improve their iSGC_MY,CR_ and iSGC_PY,CR_, thus resulting on the discussed correlations between EBVs and iSGC’s.

With respect to the relationship between MSPD and SCS, for which the mean iSGC was 0.10, both these traits were positively correlated to iSGC_MSPD,SCS_, confirming that a greater genetic potential for MSPD does lead to an increase in the genetic susceptibility to mastitis. While the 95% confidence ellipses in Figure 5 suggest that the relationship between MSPD-iSGC_MSPD,SCS_ remains stable, throughout the years of selection, the positive correlation between SCS-iSGC_MSPD,SCS_ appears to be attenuated in the more recent years.

Last, but not least, with respect to the relationship between SCS and CR, for which the mean iSGC was -0.06, only CR presented a correlation with iSGC_SCS,CR_, being this correlation negative, confirming that a greater genetic potential for CR leads to a decrease in the genetic susceptibility to mastitis. Moreover, from the 95% confidence ellipses in Figure 5, we observed that this negative correlation between SCS-iSGC_SCS,CR_ and between CR-iSGC_SCS,CR_ remains rather stable throughout the years of selection.

## Conclusion

Our study provided a number of insights with respect to the genetic correlations between traits. As proposed in our objectives, we extended the already existing inferences about how selection and random drift affect genetic variances, to how these factors affect genetic correlations, accounting for selection in a multi-trait context. To our knowledge, no comparable theoretical study has been previously performed. This theoretical front of our study revealed that, in a multi-trait selection scheme, positive genetic correlations will tend to be attenuated (*i*.*e*. decrease), and depending on the intensity of selection, they may revert to negative values. Uncorrelated traits will be driven to present a negative genetic correlation. Negatively correlated traits may be either intensified or attenuated (*i*.*e*. decrease or increase, respectively), depending on the intensity of selection and on the weights (defined by the breeding goals) used on the unified selection index. As a general rule, two traits with a small to moderate negative genetic correlation will tend to further intensify their negative genetic correlation, if the unified selection index has very different weights for these traits (*e*.*g*. 90% for one trait and 10% for the other). Moreover, in order to further our understanding of genetic correlations, we proposed an alternative method to calculate genetic correlations between traits, called by us as an individualized sire genetic correlation (iSGC), that allowed us to obtain genetic correlations at the individual level, rather than the typically estimated genetic correlations as a population parameter. The iSGC searches for individual-specific patterns of correlations, and when measured on sires through the EBVs of their daughters, it also considers the recombination of the genetic background, being the iSGC a potentially less biased measure of genetic correlations between traits. Finally, a case-study on real data was presented for the French Holstein dairy cattle population, with trajectories of genetic correlations analyzed in comparison to the theoretical results, as well as in comparison to the breeding goals imposed to this population throughout the thirty years comprised in the data set, for all the pairs of five traits of commercial interest: milk and protein yield, milking speed, somatic cell score, and cow conception rate. The trajectories of iSGC were in better agreement with the expectations from the developed theoretical calculus, enhancing our belief that this measure is indeed a less biased manner to infer genetic correlations in a breeding population.

## Declarations

### Ethics approval and consent to participate

Not applicable.

### Consent for publication

Not applicable.

### Availability of data and materials

The real data analysed are from a commercial source and are not publicly available.

## Competing interests

The authors declare that they have no competing interests.

### Funding

This project was supported by the European Union’s Horizon 2020 Programme for Research & Innovation under grant agreement n°101000226 (RUMIGEN), and by ANR PIA funding: ANR-20-IDEES-0002 (HISTOIRE).

## Authors’ contribution

BCDC worked on the theoretical derivations/results of this study, executed the analyses on the real data, conceptualized the calculus of the iSGC (individualized sire genetic correlation), and wrote the manuscript. MRM worked on the theoretical derivations/results of this study, and shared relevant discussions on both the theoretical aspects and results observed for the real data. JV conceptualized the study, and shared relevant discussions on both the theoretical aspects and results observed for the real data. NLG shared relevant discussions on the theoretical derivations of this study. FS and PC shared relevant discussions on genetic correlations that led to the concept of iSGC. DB assisted on the triage of the real data used to perform the analyses and on the definition of the calculus of the iSGC, shared relevant discussions on the Gibbs sampler method used to implement the calculus of the trajectories of genetic correlations, and on the theoretical aspects and results observed for the real data. SA assisted on the triage of the real data used to perform a preliminary analysis that inspired the final study, and shared relevant discussion on the theoretical aspects and results observed for the real data. SM conceptualized the study, and shared relevant discussions on both the theoretical aspects and results observed for the real data. All authors read and approved the final manuscript.

## Acknowledgements

Not applicable.

## Appendix A: Moments of the truncated bivariate normal distribution

Let X = [X_1_ X_2_] ∼ N(µ, Σ); E(X) = µ = [μ_1_ μ_2_] and Var 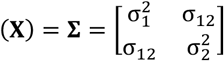. We now define the linear combination 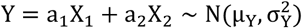, such that μ_Y_ = a_1_μ_1_ + a_2_μ_2_ and 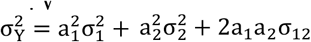. Our interest lies on the conditional distribution:

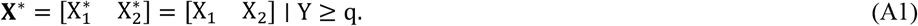

Since X = [X_1_ X_2_] ∼ N(µ, Σ), the distribution of X^∗^ = [X_1_ X_2_] ∣ Y ≥ q, is given by,

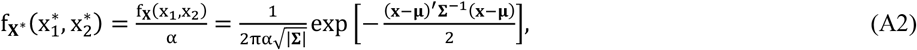

for every x_1_, x_2_ satisfying the linear truncation y = a_1_x_1_ + a_2_x_2_ ≥ q, or equivalently for every x_1_ ∈ (−∞, ∞) and 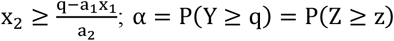, such that Z ∼ N(0,1) and 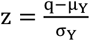. A very relevant remark is that the objective of the linear truncation imposed by us to X_1_ and X_2_ has the goal to maintain or increase their means, and therefore a_1_, a_2_ ≥ 0. This constrain on the coefficients of the linear truncation is relevant to interpret its consequences to the parameters of the distribution.

Define 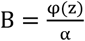 in which φ(z) is the density of the N(0,1) at the point 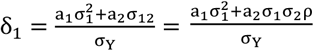, and 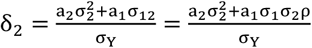, such that 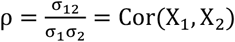. The first truncated moments are [28]:

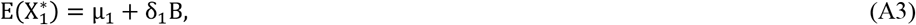

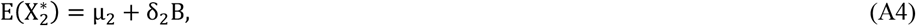

and the second truncated moments are [28]:

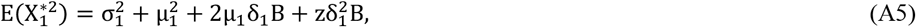

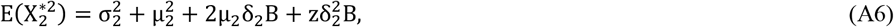

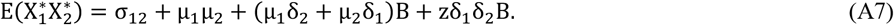

Finally, the truncated variances and covariance are:

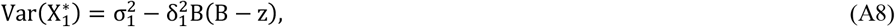

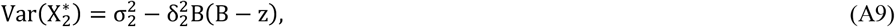

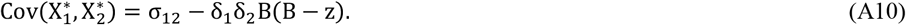

Note that B(B − z) ∈ (0,1), with 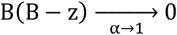 and 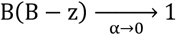, (Figure A1) and since both 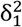 and 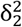 are positive, the truncation will always decrease the original variances 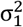 and 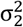.

Now, the truncated correlation is:

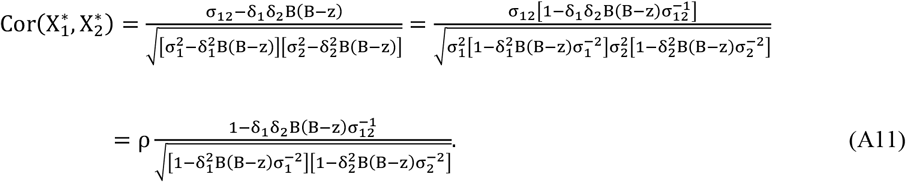

**Figure A1:**
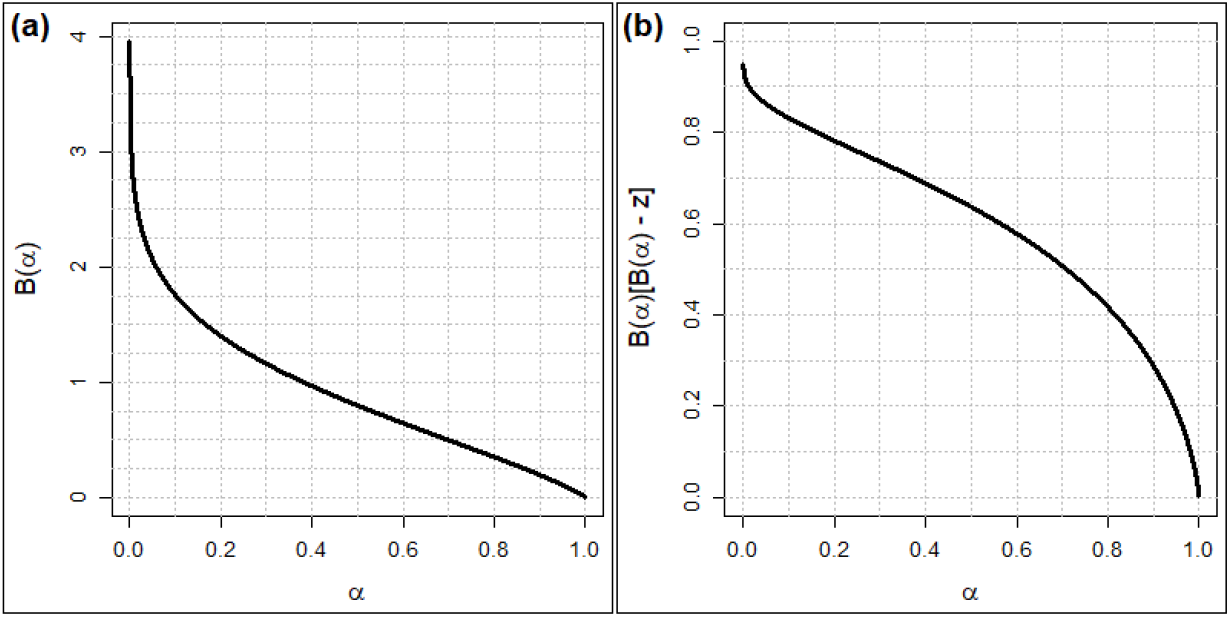
Curves of **(a)** 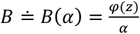, and **(b)** *B*(*B* − *z*) ≐ *B*(*α*)[*B*(*α*) − *z*], in which *z* = Φ^−1^(1 − *α*), for every *α* ∈ [0,1].

## Appendix B: Moments of a sequentially truncated bivariate normal distribution

We shall extend the moments of the truncated bivariate normal distribution presented in Appendix A, to a sequence of truncations. Let 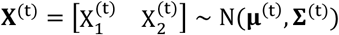 be a bivariate normal variable after the t − th truncation; 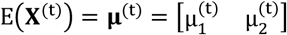 and 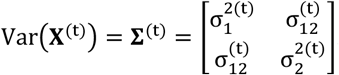. We now define the linear combination of the t − th truncation as 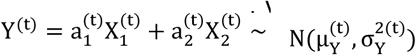, such that 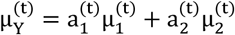 and 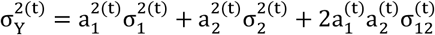.

Using the moments from the truncated bivariate normal distribution as presented in Appendix A, we have that the means and (co)variances of the α-quantile that satisfies Y^(t)^ ≥ q^(t)^ are:

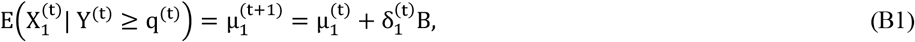

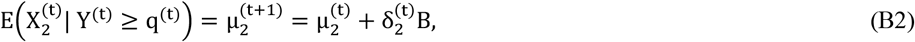

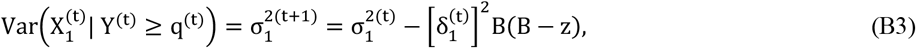

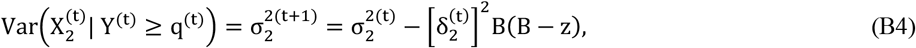

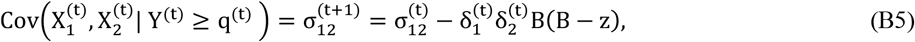

such that 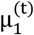 and 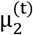 are the means at the t − th truncation; 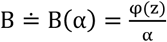 in which φ(z) is the density of Z ∼ N(0,1) at the point 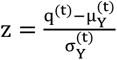, such that 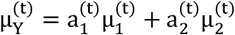 and 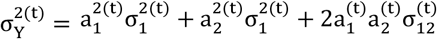, satisfying 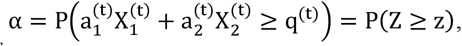, therefore 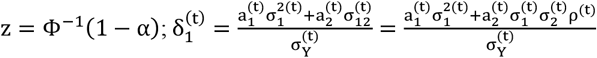 and 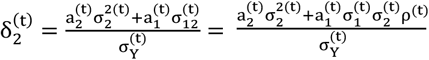.

Operating equations (B1-B5) recursively, and letting 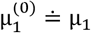 and 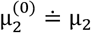 to simplify notation, we obtain,

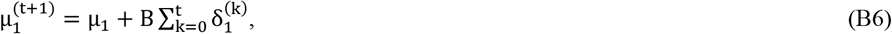

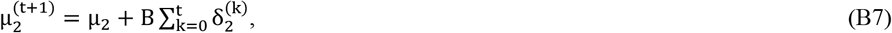

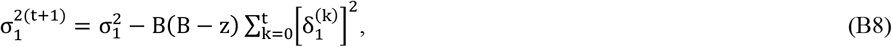

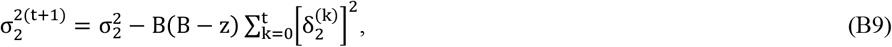

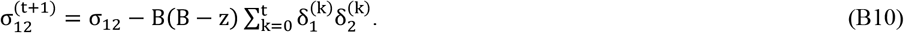

Finally, letting 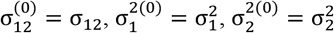, and ρ^(0)^ = ρ to simplify notation, we have

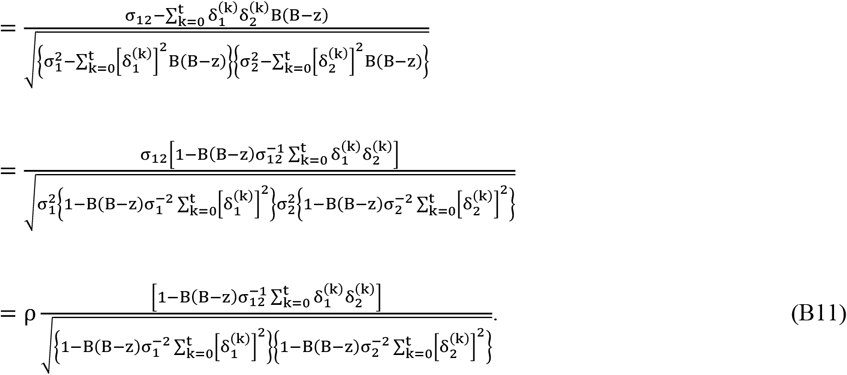

## Appendix C: Consequences of the linear truncation on the correlation parameter

By differentiating the right-hand side of equation (A11), with respect to ρ, one can verify that 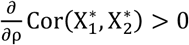, for every ρ ∈ (−1,1), indicating that 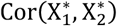 is a strictly increasing function with respect to ρ, within its parametric space. It is then sufficient to identify which value of ∈ (−1,1) satisfies 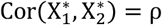, to then define when 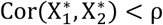 and 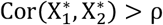. From the right-hand side of equation (A11), it can be verified that 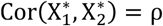, *i*.*e*. 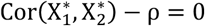, for 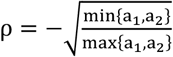, and therefore, since 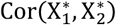 is a strictly increasing function with respect to ρ within its parametric space, we have that 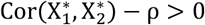 for 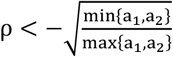, and 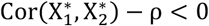 for 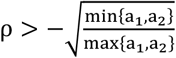. Thus, the three possible effects on the correlation due to the linear truncation are:

1. ρ decreases 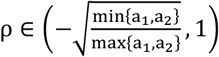
2. ρ increases if 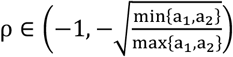
3. ρ remains unchanged if 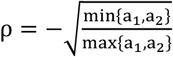.

It is important to remark that the meaning of ρ decreasing or increasing has different interpretations, according to the signal of the original ρ. If ρ > 0, decreasing means that the correlation is attenuating, and increasing means that the correlation is intensifying. Now, if ρ < 0, decreasing means that the correlation is intensifying, and increasing means that the correlation is attenuating.

Table C1 summarizes the consequences to the variance parameters when the linear truncation Y = a_1_X_1_ + a_2_X_2_ ≥ q is imposed, constrained to a_1_, a_2_ ≥ 0, assisted by the visual assessment presented in the left panels in Figure C1, which explore different combinations of a_1_, a_2_; the right panels in Figure C1 exemplify the medium-term effect after ten generations of the linear truncation (right panels) for different starting correlations (null, positive, and negative) and the different combinations of a_1_, a_2_. Note that the consequences of the linear truncations Y_1_ = a_1_X_1_ + a_2_X_2_ ≥ q and Y_2_ = a_2_X_1_ + a_1_X_2_ ≥ q are the same, given the point of inflection 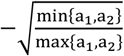 of the truncated ρ.

**Table C1:** Consequences of the linear truncation Y = a_1_X_1_ + a_2_X_2_ ≥ q constrained to a_1_, a_2_ ≥ 0, on the variances and correlation parameter of the bivariate normal distribution, for the different values of the original correlation. The consequences are summarized as an increase (↑), decrease (↓), or no changes (−--) in the parameters of interest, including the final effect on the correlation (*i*.*e*., whether it attenuates, intensifies, or becomes negative).

| Consequence of the linear truncation $a_1X_1 + a_2X_2 \geq q$ | | | |
| --- | --- | --- | --- |
| $\rho$ | $\sigma_1^2$ | $\sigma_2^2$ | $\rho$ truncated |
| $\left(-1, -\sqrt{\frac{\min\{a_1, a_2\}}{\max\{a_1, a_2\}}}\right)$ | ↓ | ↓ | ↑ (attenuates) |
| $-\sqrt{\frac{\min\{a_1, a_2\}}{\max\{a_1, a_2\}}}$ | ↓ | ↓ | --- |
| $\left(-\sqrt{\frac{\min\{a_1, a_2\}}{\max\{a_1, a_2\}}}, 0\right)$ | ↓ | ↓ | ↓ (intensifies) |
| 0 | ↓ | ↓ | ↓ (becomes neg.) |
| (0,1) | ↓ | ↓ | ↓ (attenuates) |

**Figure C1:**
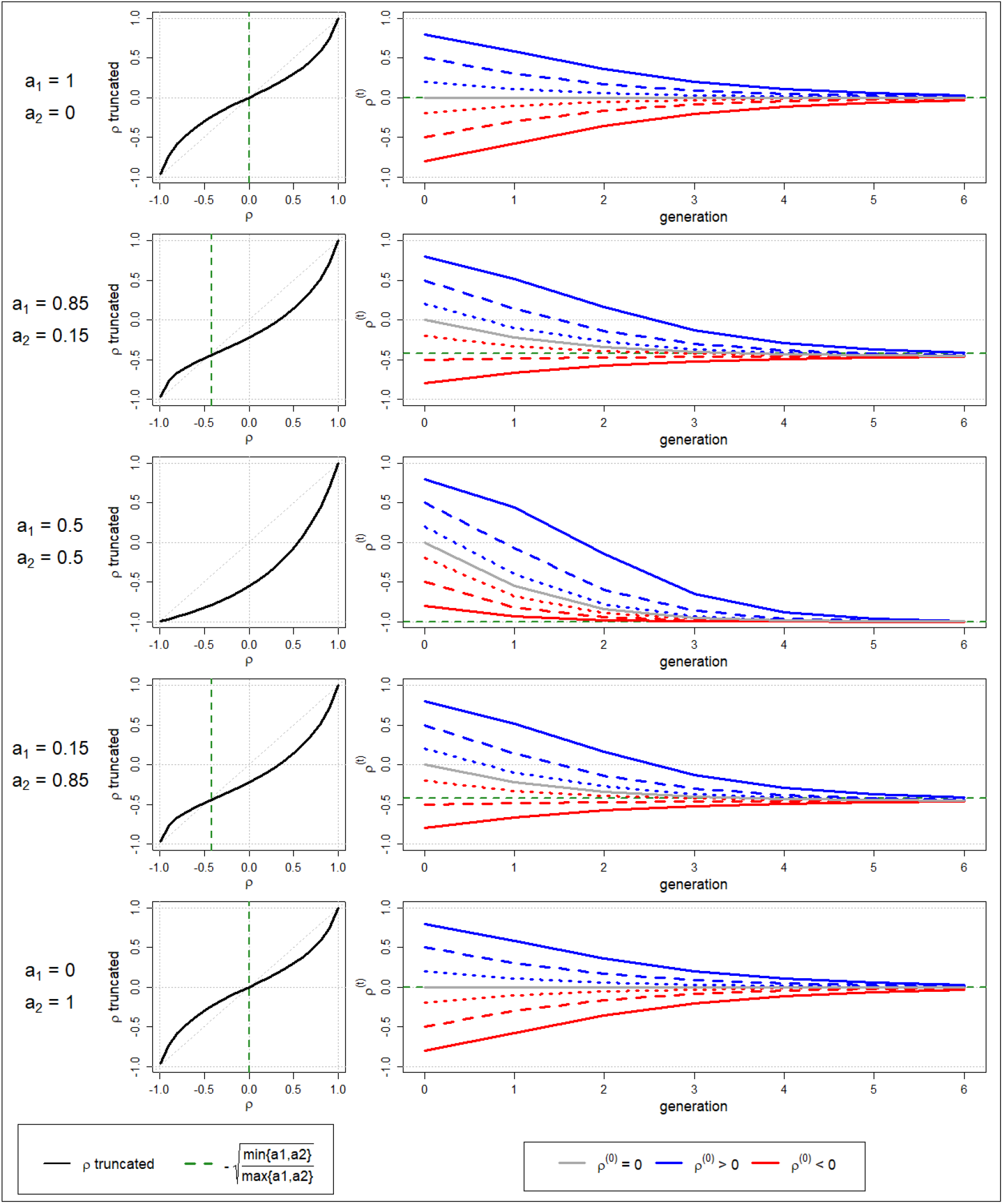
Illustration of the consequences of the linear truncation Y = *a*_1_*X*_1_ + *a*_2_*X*_2_ ≥ *q* on the correlation parameter of the bivariate normal distribution, for *P*(Y ≥ *q*) = 0.35 and for different combinations of *a*_1_, *a*_2_, from the functional analysis perspective (left panels), and the medium-term effect after ten sequential truncations for different starting correlations, either null, positive, or negative (right panels). Incomplete lines before the tenth sequential truncation occurred when variances reached zero.

## Notes

### Competing Interest Statement

The authors have declared no competing interest.

